# Aesthetic and physiological effects of naturalistic multimodal music listening

**DOI:** 10.1101/2022.07.02.498390

**Authors:** Anna Czepiel, Lauren K. Fink, Christoph Seibert, Mathias Scharinger, Sonja A. Kotz

## Abstract

Compared to audio only (AO) conditions, audiovisual (AV) information can enhance the aesthetic experience of a music performance. However, such beneficial multimodal effects have yet to be studied in naturalistic music performance settings. Further, peripheral physiological correlates of aesthetic experiences are not well-understood. Here, participants were invited to a concert hall for piano performances of Bach, Messiaen, and Beethoven, which were presented in two conditions: AV and AO. They rated their aesthetic experience (AE) after each piece (Experiment 1 and 2), while peripheral signals (cardiorespiratory measures, skin conductance, and facial muscle activity) were continuously measured (Experiment 2). Factor scores of AE were significantly higher in the AV condition in both experiments. LF/HF ratio, a heart rhythm that represents activation of the sympathetic nervous system, was higher in the AO condition, suggesting increased arousal, likely caused by less predictable sound onsets in the AO condition. We present partial evidence that breathing was faster and facial muscle activity was higher in the AV condition, suggesting that observing a performer’s movements likely enhances motor mimicry in these more voluntary peripheral measures. Further, zygomaticus (‘smiling’) muscle activity was a significant predictor of AE. Thus, we suggest physiological measures are related to AE, but at different levels: the more involuntary measures (i.e., heart rhythms) may reflect more sensory aspects, while the more voluntary measures (i.e., muscular control of breathing and facial responses) may reflect the liking aspect of an AE. In summary, we replicate and extend previous findings that AV information enhances AE in a naturalistic music performance setting. We further show that a combination of self-report and peripheral measures benefit a meaningful assessment of AE in naturalistic music performance settings.

## 1 Introduction

There is a clear consensus that listening to music induces aesthetic experiences, with humans augmenting such experiences by optimising the ‘where’ and ‘how’ we listen to music, such as in concerts (Sloboda et al., 2012; Sloboda & O’Neill, 2001; Wald-Fuhrmann et al., 2021). Although the aesthetic experience (AE) of music is enhanced in a concert by several aspects (see Wald-Fuhrman et al., 2021 for an overview), one explored here is visual information. While previous work showed that visual cues enhance self-reported musical evaluation of music performances (e.g., see Platz & Kopiez, 2012 for a meta-analysis), some gaps in the literature remain. Firstly, most studies comparing audiovisual (AV) and audio only (AO) musical performances have been conducted in laboratory settings; to test a more genuine AE, it is imperative to use a more naturalistic situation. Secondly, only two studies so far explored physiological responses between AV and AO musical performances (Chapados & Levitin, 2008; Vuoskoski et al., 2016), but their findings are contrary to each other. Thus, the current study aimed to specify the link between modality (AO vs. AV), AE, and peripheral physiological responses in a naturalistic music performance setting, i.e., a piano concert.

While the initial study of AE has had a strong philosophical focus, AE is currently of great interest in cognitive neuroscience and the neuroscientific subdiscipline of neuroaesthetics. Here, perception, emotion, and appreciation are considered to influence AE (for comprehensive reviews, see Anglada-Tort & Skov, 2020; Brattico & Pearce, 2013; Juslin, 2013; Pelowski et al., 2016; Schindler et al., 2017). Specific to the dynamic nature of music, Brattico and colleagues (2013) proposed that the AE of music listening is composed of a chronometry of components: 1) perceptual sensory processes (feature analysis/integration) as well as early emotional reactions (e.g., startle reflex and arousal), 2) cognitive processes (based on long-term knowledge, such as harmonic expectancy), and 3) affective processing (including perceived and felt emotions). A combination of these processes that involve somatomotor processes interacting with the listener themselves (in terms of cultural knowledge, musical expertise, etc.) and external context (e.g., social setting), result in 4) aesthetic responses (emotions, judgements, and liking). Brattico et al. (2013) presented neurophysiological correlates that might reflect these processes. Namely, sensory processes should be reflected in early event-related potentials (ERPs) and in early auditory processing areas (sensory cortices, brainstem). More cognitive (‘error’ and ‘surprise’) components should be reflected in the MMN and P300 and non-primary sensory cortices. Finally, (aesthetic) emotion and judgements should be reflected in the late potential component (LPC) and reward and emotion areas in the brain. Research further suggests that (synchronisation of) certain brain oscillations are related to music-evoked pleasure, particularly frontal theta oscillations (Ara & Marco-Pallarés, 2020; Sammler et al., 2007; Tervaniemi et al., 2021), parieto-occipital alpha (Nemati et al., 2019), theta (Chabin et al., 2020) and theta phase synchronisation (Ara & Marco-Pallarés, 2020, 2021), as well as the inter-brain synchrony (IBS) of frontal and temporal theta in shared musical pleasure (Chabin et al., 2022).

Although some work has explored music-evoked pleasure with EEG in the more naturalistic setting of a concert hall (Chabin et al., 2022), measuring brain activity in such settings comes with significant challenges. A more accessible approach, however, has been to measure peripheral physiological responses in naturalistic settings such as theatres (Ardizzi et al., 2020), concert halls (Egermann et al., 2013), and cathedrals (Bernardi et al., 2017). Peripheral measures include the somatic (voluntary muscle) and autonomic nervous systems (ANS), of which the latter comprises the sympathetic (‘fight-or-flight’) and parasympathetic (‘rest-and-digest’) nervous systems (SNS, PNS). In naturalistic settings, previous work revealed (synchronised) physiological arousal responses in audiences occur in relation to surprising, emotional, and structural moments in music such as transitional passages, boundaries, and phrase repetitions (Czepiel et al., 2021; Egermann et al., 2013; Merrill et al., 2021). Such peripheral measures are likewise mentioned in the AE chronometry approach (Brattico et al., 2013) as reflecting tension and chill responses (Grewe et al., 2009; Salimpoor et al., 2009). However, unlike brain regions (fMRI) and the latency/polarity of (EEG/MEG) components, that can be attributed to psychological processes (Kappenman & Luck, 2011), peripheral responses are mainly characterised according to increased/decreased activity, making it more difficult to separate responses relating to distinct sensory, cognitive, and/or aesthetic processes. Thus, rather than taking a superficial understanding that such measures directly index a pleasurable experience, a more thorough biological understanding is required to appropriately interpret the meaning of such measures (see e.g., Fink et al., 2023, for an example in pupillometry).

The current dependent measures of interest, which have also previously been used in research on musical aesthetics (e.g., Grewe et al., 2009; Salimpoor et al., 2009), range from involuntary ANS responses to voluntary motoric control, namely: skin conductance, heart, respiratory, and muscle activity. Skin conductance (SC, also known as electrodermal activity, EDA) measures activation of sweat glands, which are innervated by the SNS only. The heart consists of cardiac muscle (involuntary control), with SNS (via sympathetic nerves) and PNS (vagus) innervations that increase and decrease heart rate (HR), respectively. Typically, HR fluctuates and is measured by different heart rate variability (HRV) measures. These measures can be in the time-domain, for example, the standard deviation between interbeat intervals, or in the frequency-domain, for example, power of certain frequency bands related to SNS and PNS activation. Power at a high frequency (HF, 0.4-0.15Hz) component is attributed to PNS activity, while power at a low frequency (LF, 0.04 - 0.15 Hz) component seems to reflect both PNS and SNS influences; thus, the LF/HF ratio is used to represent SNS activity (Malik, 1996; Shaffer & Ginsberg, 2017). Respiratory activity encompasses both involuntary control - where the lungs are innervated by both SNS and PNS, which dilate and constrict the bronchioles, respectively as well as voluntary control (Purves & Williams, 2001). The somatic (muscle) system consists mainly of skeletal (voluntary) muscle; commonly measured are the facial muscles of zygomaticus major (‘smiling’) and corrugator supercilii (‘frowning’). Although under voluntary control, certain facial muscle responses may be partly unconscious (i.e., occur without attention or conscious awareness, Dimberg et al., 2000). Overall, SC, heart, respiration, and facial muscle activity broadly relate to arousal and valence^1^. Higher arousal has been associated with SNS activation, such as increased sweat secretion, increased LF/HF ratio, HR and RR acceleration, and decreased HF power (Di Bernardi Luft & Bhattacharya, 2015; Shaffer & Ginsberg, 2017), while zygomatic and corrugator muscle activity seem to reflect positive and negative valence, respectively (Bradley & Lang, 2000; Cacioppo et al., 2000; Dimberg et al., 2000; Lang et al., 1993; Larsen et al., 2003, though see discussion below).

Although broadly reflecting arousal and valence, peripheral measures have been related to sensory, cognitive, and aesthetic experiences with regard to acoustic/musical stimuli in separate studies. Increased SC and HR patterns have been related to early sensory reactions to an acoustic signal referred to as an orienting response/startle reflex (Barry, 1975; Barry & Sokolov, 1993; Graham & Clifton, 1966; Roy et al., 2009). Physiological changes occur in response to cognitive music processes such as recognising unexpected harmonic chords (Koelsch et al., 2008; Steinbeis et al., 2006) and deviant stimuli (in an MMN-like paradigm, Chuen et al., 2016; though see Lyytinen et al., 1992), which might be enhanced by attention (Frith & Allen, 1983). In more naturalistic music listening, many studies showed that arousing music (faster tempi and unpredictable harmony) increase SC, HR, and RR (Bernardi et al., 2006; Coutinho & Cangelosi, 2011; Czepiel et al., 2021; Dillman Carpentier & Potter, 2007; Egermann et al., 2013, 2015; Khalfa et al., 2002; Krumhansl, 1997), though we note this result is not consistent across studies, for reviews see (Bartlett, 1996; Hodges, 2009; Koelsch & Jäncke, 2015). In terms of valence, researchers have shown that zygomaticus activity increases during happy music (Lundqvist et al., 2008). However, other work showed it can increase during unpleasant (dissonant) music (Dellacherie et al., 2011; Merrill et al., 2021). This conflict suggests that perhaps the activation of the smiling muscle is not just related to valence (see also Wingenbach et al., 2020). Peripheral responses have likewise been related to aesthetic experience of music, or least music-evoked “chills” (frissons), which increases SC, HR, RR and EMG (Blood & Zatorre, 2001; Craig, 2005; Grewe et al., 2009; Salimpoor et al., 2009). Hence, evidence suggests that peripheral measures can reflect (a mixture of) the sensory, cognitive and/or preference parts of the AE, rather than being a direct index of AE. Therefore, it is of importance to collect self-report measures to further interpret the peripheral responses to AV and AO comparisons.

In terms of modality effects on self-reports, audio information seems to be consistently influenced by performer movement. In one percussion study, pairing visual gestures that created long notes to acoustic sounds of short notes resulted in short sounds being perceived as longer sounding notes (Schutz & Lipscomb, 2007); an effect later shown to be consistent in percussive (but not sustained) sounds when the sound appears after a gesture (Schutz & Kubovy, 2009). In piano performances, one acoustic performance was paired with four videos: one as the original performance and three pianist ‘doubles’. Ninety-two out of ninety-three participants perceived differences between the performances, although the sound remained identical (Behne & Wöllner, 2011). With regard to more aesthetic influences, several studies that compared uni- and bimodal versions of music performances showed visual cues enhance a listener’s perception of performance quality (Waddell & Williamon, 2017), musical expertise (Griffiths & Reay, 2018; Tsay, 2013), musical expression (Broughton & Stevens, 2009; Davidson, 1993; Lange et al., 2022; Luck et al., 2010; Morrison & Selvey, 2014; Vines et al., 2011; Vuoskoski et al., 2014), perception of emotional intention (Dahl & Friberg, 2007; Vines et al., 2006), and felt emotion (Van Zijl & Luck, 2013). As AE is related to the appreciation of performance expressiveness, quality, and emotion (Brattico & Pearce, 2013; Juslin, 2013), this research, as well as a meta-analysis (Platz & Kopiez, 2012), showed that AE increases with additional visual cues. One neuroaesthetic theory that could further explain this enhanced AE postulates that visual information may increase embodied simulation, which subsequently increases AE (Freedberg & Gallese, 2007; Gallese & Freedberg, 2007). Support for this idea comes from studies showing higher activation in the action observation network when viewing movements that are rated as aesthetically pleasing (Cross, 2011).

However, this enhanced AE effect has been mostly assessed in laboratory settings. Recent studies are increasingly exploring such experiences in live concerts (Chabin et al., 2022; Coutinho & Scherer, 2017; Czepiel et al., 2021; Scherer et al., 2019; Swarbrick et al., 2019; Tervaniemi et al., 2021), where participants report experiencing stronger emotions (Gabrielsson & Wik, 2003; Lamont, 2011); however, Belfi et al. (2021) found that felt pleasure did not differ between live and an audiovisual recording of the same performance. Focusing more specifically on the role of modality, to date only a few studies compare responses to AV vs. AO conditions in naturalistic settings. Compared to eyes-closed conditions, eyes-open conditions increased movement energy and interpersonal coordination, suggesting that visual information may enhance the social aspect of live pop/soul music (Dotov & Trainor, 2021). Coutinho & Scherer (2017) compared emotional responses in a live AV performance to recorded AV, AO, and VO performances of Schubert Lieder, where the live AV condition had significantly higher wonder and significantly lower boredom ratings. Although these two studies highlight the difference between genres and the affordances that visual information can give (focus on seeing other audience members/musicians in popular/classical music, respectively), they essentially show that additional information enhances the (social/emotional) experience. We stress that it is not trivial to replicate findings from the lab to a more naturalistic setting, since, for example, well documented effects of familiarity and body movement on music appreciation found from lab studies were not replicated in a field study (Anglada-Tort et al., 2019). It is also worth extending Coutinho & Scherer (2017), since they focus on the more emotional part of AE, and only collected data from an AV modality in a naturalistic setting (other modalities were tested in a lab-like setting). The current study thus compares modalities in one naturalist setting to examine more specifically the judgement and preference components of AE.

Two previous studies have compared peripheral physiological responses as a function of modality during music performances and serve as the starting point for the current work. Chapados & Levitin (2008) found that self-reported tension as well as SC were both highest in AV conditions. However, Vuoskoski et al. (2016) found that, although self-reported intensity, high energy arousal, and tension were highest in AV conditions, SC was actually highest in AO conditions. While the discrepancy between these two studies could relate to the different styles and instruments used (which offer different expressive affordances), Vuoskoski et al. (2016) argued that SC might be higher during AO performances due to musical expectancy (Huron, 2006; Juslin & Västfjäll, 2008). More specifically, as visual information increases listeners’ ability to predict upcoming musical events, AV stimuli are less surprising. Indeed, this idea is supported by speech studies focusing on the N100, an EEG event-related potential component that reflects early sensory processing, where a larger N100 amplitude can indicate a response to a less predictable stimulus. The N100 component is enhanced in AO (compared to AV) conditions in speech (Klucharev et al., 2003; van Wassenhove et al., 2005), emotional expression (Jessen & Kotz, 2011), as well as non-speech events such as clapping (Stekelenburg & Vroomen, 2007). These findings corroborate the idea that the lack of visual information makes sound onsets less predictable.

Together, this evidence suggests that peripheral responses might be 1) higher in AO conditions if they reflect sensory processing, or 2) higher in AV conditions if they reflect the enhanced emotional and/or appreciation aspects of AE. If peripheral physiological responses reflect sensory processing, we would expect to replicate results from Vuoskoski et al. (2016) and find increased physiological activity in AO conditions. However, if physiological responses reflect the more emotional/aesthetic aspects, we would expect to replicate results from Chapados & Levitin (2008) and find increased physiological responses in AV conditions.

In summary, more research is needed to assess modality effects that enhance aesthetic experience in a more naturalistic setting. Further, the peripheral physiological correlates of aesthetic effects are so far inconsistent. The current study consists of two experiments that examine AE and physiology between AV and AO conditions in a concert hall setting. In both Experiments, we recorded behavioural responses and tested the hypothesis that AE will be higher in the AV condition. In Experiment 2, we additionally collected physiological responses and tested the hypothesis put forward by Vuoskoski et al. (2016) that peripheral physiological activity should be higher in AO conditions.

## 2 General Method

### 2.1 Overview

We present two experiments, each consisting of two concerts. Experiment 1 (Concerts 1 and 2) measured behavioural ratings, while Experiment 2 (Concerts 3 and 4) measured both behavioural ratings and physiological responses. Both involve the same stimuli and the same within-subjects experimental design: participants listening to piano performances of Bach, Beethoven, and Messiaen, in AO and AV conditions. Modality order was counterbalanced across concerts.

### 2.2 Stimuli

Upon engaging a pianist, three musical pieces were selected from their repertoire in accordance with the pianist and musical experts to represent various emotional expressions (cheerful, sad, and ambiguous) and musical styles (Baroque, Classical-Romantic, and 20^th^ century music): Johann Sebastian Bach: Prelude and Fugue in D major (Book Two from the Well-Tempered Clavier, BWV 874), Ludwig Van Beethoven: Sonata No. 7, Op. 10, No. 3, second movement (Largo e mesto), and Olivier Messiaen: *Regard de l’Esprit de joie* (No. 10 from Vingt Regards sur L’Enfant-Jésus). These pieces were presented to the participants during each concert twice in the two different modalities: in audiovisual (AV) and an audio only (AO) versions. We considered this repetition of pieces as a naturalistic part of the design as piece repetition is a practice (although not extremely common) in concert programming (Halpern et al., 2017).

Both AV and AO presentations of the music pieces were performed by the same pianist, playing on the same piano (Steinway B-211), in the same concert hall. AV versions of the music pieces were performed live during the concerts and the audience could see and hear the pianist performing the music. AO versions of the music pieces were recorded in the same concert hall, on the same piano in advance of the concerts, without an audience. The AO versions were presented during the concerts via a stereo setup with high-quality full-range loudspeakers (Fohhn LX-150 + Fohhn XS-22), so that the audience could only hear the music. During this time, the pianist was backstage, so that the audience could only see the piano. The playback AO versions were the same in all concerts in both experiments. To ensure similarity of sound levels between AO and AV presentations, a trained sound engineer checked that the loudness across the modalities was equal.

Although modality conditions were controlled as much as possible, we would assume that repeated performances of the same musical piece might have slight deviations from each other, even when performed by a highly trained professional musician (Chaffin et al., 2007). Therefore, we checked that the stimuli nonetheless were comparable enough to eliminate confounding variables of potential acoustic differences between AV versions (different for each concert) and AO versions (the same across all concerts). We differentiated between score-based features and performance-based features (Goodchild et al., 2019). The former refers to features that come from the notated scores (e.g., harmonies), which should remain the same across performances (assuming no errors in playing the scores). The latter refers to features that may also be notated in the scores (e.g., dynamic markings) but might deviate more depending on the performances, such as tempo, loudness, and timbre. Tempo was extracted using a combination of MIDI information for each note and manually locating the beat (using Sonic Visualiser, Cannam et al., 2010), where inter-beat intervals were obtained to calculate continuous beats per minute (bpm). Loudness and timbre were extracted from the audio signal using MIRToolbox (Lartillot & Toiviainen, 2007) in MATLAB, with RMS (*mirrms*) and spectral centroid (*mircentroid*) representing loudness and timbre, respectively. In checking multicollinearity (Lange & Frieler, 2018), none of the features correlated highly, confirming that each feature represented an independent aspect of the music. Each of the features were averaged into average bins per bar (American: measure) to account for slight timing deviances between performances. The features over time were very similar (see Supplementary Figures 1-6 in Supplementary Materials). This similarity was confirmed by significant correlations between concerts, all with *r* values > 6 (see Supplementary Materials, Supplementary Table 1), suggesting that all performances were acoustically comparable.

### 2.3 Questionnaires

Questionnaires were presented after each musical piece to assess three types of questions. Firstly, we assessed the ‘naturalness’ of the concert by asking to what extent the experimental components of the setting (e.g., measurement of the behavioural responses) disturbed the concert experience, where ‘disturbed by measurement’ was rated from 1 (strongly disagree) to 7 (very much agree). We further assessed familiarity with the style of music as well as whether the participant knew the specific piece of music. This was rated from 1 (not at all familiar) to 7 (very familiar). Thirdly, we assessed the main dependent variable of interest: aesthetic experience (AE). As an AE is made up of several components (Brattico & Pearce, 2013; Schindler et al., 2017), we assessed the aesthetic experience with a set of eight individual items, consisting of how much they liked the piece, how much they liked the interpretation of the piece, and how absorbed they felt in the music (see Supplementary Materials for all questions).

### 2.4 Procedure

Participants were invited to attend piano concerts that took place at the ArtLab of the Max-Planck-Institute for Empirical Aesthetics in Frankfurt, a custom-built concert hall seating 46 audience members (https://www.aesthetics.mpg.de/en/artlab/information.html). Concerts were kept as identical as possible for factors such as lighting, temperature, and timing. Prior to the concert, participants were informed about the experiment and filled in consent forms before being seated in the ArtLab. During the concert and after each piece of music, participants answered the short questionnaire described above. All participants saw the three pieces in both conditions. For one concert per Experiment, the three music pieces were presented first in the AO modality, and then repeated in the AV modality. Modality order was counterbalanced so that in the other concert per Experiment, music pieces were first presented in the AV modality, and then again in the AO modality. An overview of the procedure and modality condition orders can be found in Figure 1. Behavioural measures were recorded in both Experiment 1 and 2. In Experiment 2 only, physiological data were additionally collected (details in Section 4.1.2 Experiment 2 Procedure).

**Figure 1.**
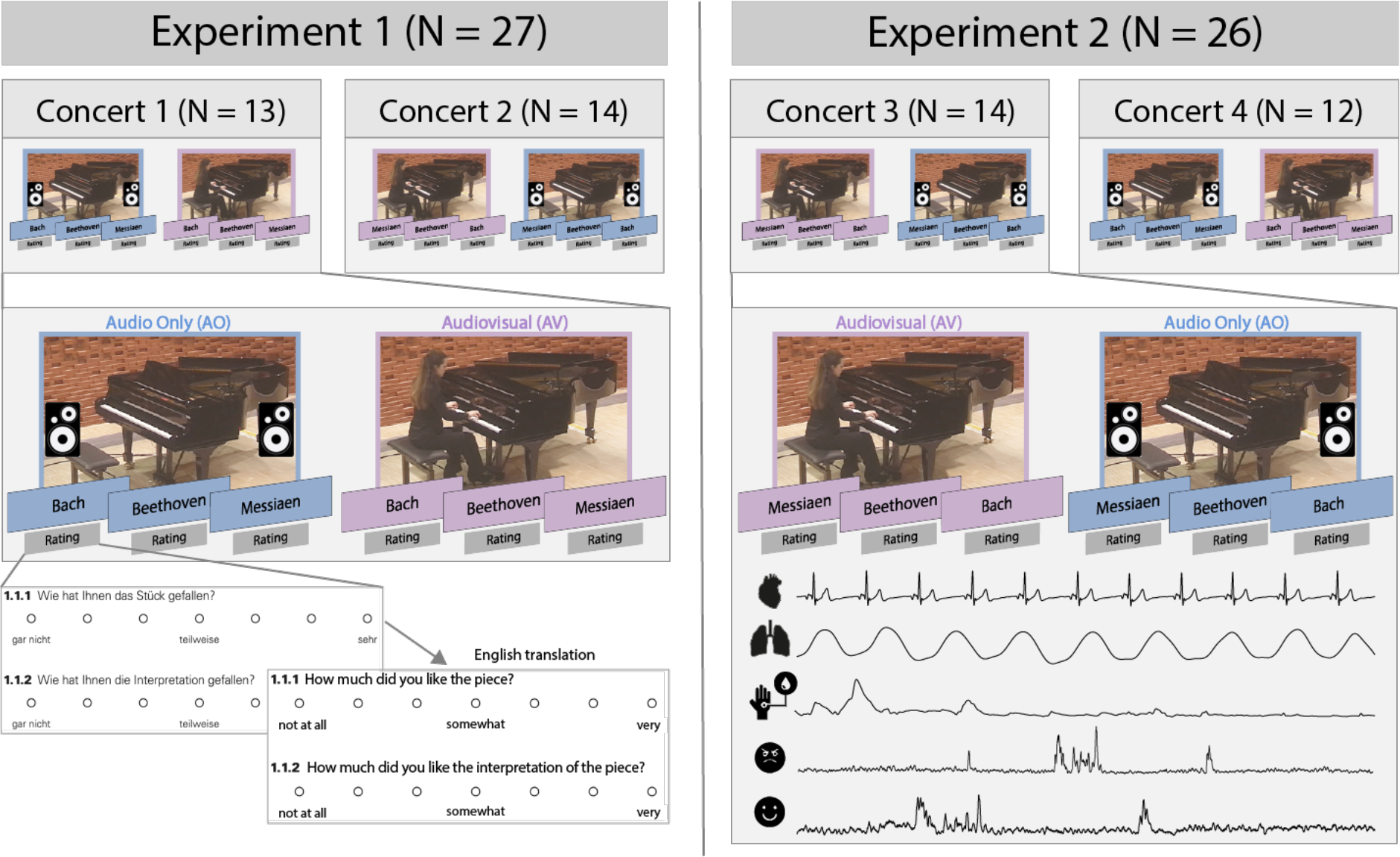
Outline of the experimental procedure in Experiment 1 (behavioural audience ratings) and Experiment 2 (audience ratings and peripheral physiological measures). Pieces were presented both in an AV version (purple boxes) and an AO version (presented via speakers, blue boxes).

### 2.5 Analysis

Statistical analyses were conducted in R and R studio (R Core Team, 2021; RStudio Team, 2021).

Items chosen for the questionnaire (see Supplementary Materials) reflect elements of an aesthetic experience. Thus, it was assumed that the items might be related to each other. Indeed, in both Experiments, items in the self-reports capturing the aesthetic experience were highly correlated. Therefore, rather than comparing modality differences for each item, we reduced the questionnaire items to an overall, more interpretable factor - that retains important information from each item - using a factor analysis (Fabrigar et al., 1999). This reduced factor yielded new factor scores that mixed scores from the original items together based on loadings, i.e., regression weights (using *fa* from the psych package, see accompanying code; Revelle, 2022). The more one item contributed to - or loaded onto - the reduced factor, the higher the ‘item loading’ was for that factor. Table 1 shows the item loadings of factors in both experiments. These factor scores were used as a new overall variable that represents a summary of the questionnaire items. Details about each factor analysis (FA) for each experiment are explained below in the experiment-specific methods.

**Table 1.** FA loadings from questionnaire items in both Experiment 1 and 2. Factor 1 for both Experiment 1 and 2 is interpreted as ‘Aesthetic experience’.

|  | Experiment 1 | Experiment 2 |
| --- | --- | --- |
| Item | Factor 1 | Factor 1 |
| Liking | 0.78 | .87 |
| Liking of interpretation | 0.69 | 0.63 |
| Absorption | 0.90 | 0.67 |
| Passive reception | 0.73 | 0.09 |
| Connection to musicians | 0.56 | 0.43 |
| Urge to move | 0.17 | 0.21 |
| Connection to co-listeners | 0.28 | 0.14 |
| Understanding | 0.27 | 0.34 |

Linear mixed models (LMMs) were run with the factor scores extracted from the factor analysis as the dependent variable, with modality (AV / AO) as a predictor (fixed effect). We also ran LMMs for each physiological measure, where modality was the predictor (fixed effect) as well as a LMM assessing relationship between factor scores and physiological measures. LMMs are more appropriate than repeated measures ANOVA, as they are more fitting for physiological data, can account for missing trials, and can model random sources of variance and non-independence in the observations (Barr et al., 2013; Page-Gould, 2016; Winter, 2013). Ratings and physiological measures were recorded multiple times from each participant, who heard the same music piece more than once, in groups for each concert. To account for these random sources of non-independence, we added random intercepts for concert, piece, and participant. Participants were nested within concerts, while participant and piece were considered crossed effects. For the physiological data, piece sections were further nested within pieces to account for observations taken within pieces (see Methods for Experiment 2). We also included a random slope for participants. Thus, the models represent the maximal random effects structure justified by the design (Arnqvist, 2020; Barr, 2021; Barr et al., 2013). While LMMs do not rely on normally distributed data, we checked linearity, homoscedasticity, and normality of residuals of the models (Winter, 2013). We also checked for model errors. All maximal models generated singular fit errors, suggesting that the model might be too complicated and/or one or more random effects have (near to) zero variance or (near-)perfect correlations. Therefore, we followed the recommended procedure of simplifying models until error is removed (Barr, 2021), ultimately selecting a model with a random effect structure that is supported by the data (Barr et al., 2013; Matuschek et al., 2017). As error-free models are generally preferred (Barr et al., 2013), we report the models that generated no errors, but report all maximal and simplified models in the Supplementary Materials. LMMs were run using *lmer* from the *lme4* packages (Bates et al., 2015; Kuznetsova et al., 2017). Significance values, effect sizes, and Akaike information criterion (AIC) were obtained from the *tab_model* function from *sjPlot* package (Lüdecke, 2023). Pairwise comparisons were run with the *emmeans* function from *emmeans* package (Lenth, 2021) with Bonferroni corrections. As a sanity check for the linear mixed models, we also ran ANOVAs (Arnqvist, 2019). Corresponding code and required to run these analyses are available at Open Science Framework (OSF) (Please note this repository is currently private and only available with this link while the manuscript is under review; it will be made public when the manuscript is accepted).

## 3 Experiment 1

### 3.1 Method

#### 3.1.1 Participants

The study was approved by the Ethics Council of the Max Planck Society and in accordance with the declarations of Helsinki. Participants gave their written informed consent. Twenty-seven participants attended the experimental concerts (13 and 14 participants in Concert 1 and 2, respectively), 18 females (9 males), with mean age of 57.96 years (SD = 20.09), who on average had 6.99 years of music lessons (SD = 7.87) and attended approximately 13 concerts in the last 12 months (M = 12.62; SD = 13.37). Participants also provided ratings on their perception being a musician (from 1 = does not apply, to 7 completely applies), most participants selected 1 (N = 13) or 2 (N = 4), and less selected 3 (N = 1), 4 (N= 2), 5 (N = 3), 6 (N = 2) and 7 (N = 2). Most had a college/university degree (N = 22), the others either vocational training (N = 2) or completed A-levels/high school (N = 3). Wilcoxon tests showed that participants did not differ in Concert 1 and 2 in terms of age (*p* = .590), musician level (*p* = .877), years of music lessons (*p* = 1.00), and number of concerts attended in the last 12 months (*p* = .173).

#### 3.1.2 Factor analysis and statistical analysis

Questionnaire items were chosen to reflect elements of an aesthetic experience. As they were highly correlated (see accompanying code), we chose to reduce these variables to an interpretable factor using factor analysis. A Kaiser-Meyer-Olkin (KMO) measure verified sampling adequacy (KMO = .801, well over the .5 minimum required) and all KMO values for individual items were > .670. Bartlett’s test of sphericity was significant, revealing that correlations between items were large enough for a FA, *X*2(28) = 408.844, *p* < .001. Kaiser’s criterion of eigenvalues > 1 and a scree plot indicated a solution with one factor. Thus, a maximum-likelihood factor analysis was conducted with one factor, which explained 37% of the variance. We took the scores of this factor and created a new variable. As items of liking, liking of interpretation, and absorption loaded highly onto this factor, and these aspects have been identified as critical aspects of an aesthetic experience (Brattico & Pearce, 2013; Orlandi et al., 2020), we referred to this new variable as the overall ‘aesthetic experience’ (AE). Nine trials with an outlier exceeding ±3 Median Absolute Deviations (MAD, Leys et al., 2013) was removed from further analyses. In total, we had 153 observations for the AE scores [(27 participants x 3 pieces x 2 modality conditions) - 9]. We compared AE factor scores between modality conditions using LMMs (see General Methods, corresponding code).

### 3.2 Results

#### 3.2.1 Assessing naturalistic situations

Results of whether the measurements disturbed the concert are shown in Table 2. The mean rating was 1.537 (SD = 1.016) out of 7, with 88% of ratings at 1 or 2 on the scale (i.e., strongly disagree or disagree that measurements disrupted the concerts, respectively). Thus, behavioural measurements did not disrupt the concert, confirming the ecological validity of the experimental setting.

**Table 2.** Ratings of feeling disturbed by the measurement, and familiarity with style and specific piece in Experiment 1.

| Ratings of feeling disturbed by the measurement |  |  |  |  |  |  |  |
| --- | --- | --- | --- | --- | --- | --- | --- |
| Rating | 1 | 2 | 3 | 4 | 5 | 6 | 7 |
|  | 69% | 19% | 5% | 3% | 3% | 1% | 0% |
| Familiarity with style of piece |  |  |  |  |  |  |  |
| Rating | 1 | 2 | 3 | 4 | 5 | 6 | 7 |
| Bach, | 0% | 2% | 4% | 11% | 18% | 26% | 39% |
| Beethoven, | 0% | 2% | 4% | 15% | 16% | 35% | 28% |
| Messiaen, | 26% | 17% | 9% | 20% | 9% | 11% | 8% |
| Familiarity with specific piece |  |  |  |  |  |  |  |
|  | 0 (No) |  | 1 (Yes) |  |  | Not sure |  |
| Bach, | 78% |  | 18% |  |  | 4% |  |
| Beethoven, | 70% |  | 26% |  |  | 4% |  |
| Messiaen, | 85% |  | 11% |  |  | 4% |  |

#### 3.2.2 Piece familiarity

Ratings for familiarity of style were similarly high for Bach (M = 5.796, SD = 1.279) and Beethoven (M = 5.630, SD = 1.248), but lower for Messiaen (M = 3.333, SD = 1.981). Most participants did not know the pieces specifically, though 18%, 26%, and 11% of participants knew the Bach, Beethoven, and Messiaen pieces, respectively.

#### 3.2.3 Aesthetic experience: Modality differences

LMMs showed modality was a significant predictor of AE (see Table 4). AV scores were significantly higher (M = 0.186, SE = .296, 95% CI [-0.962 1.33]) than AO scores (M = -0.102, SE = .297, 95% CI [-1.245, 1.04]), *t*(124)= -.240, *p* = .018) (see Figure 2). This effect was confirmed by the maximal model, despite generating a singular fit error: it yielded the same estimates and had similar effect sizes, AIC, and significance (see Supplementary Table 3). The modality effect was confirmed by an ANOVA, which yielded a significant main effect of modality (*F*(1,26) = 5.564, *p* = .026).

**Figure 2.**
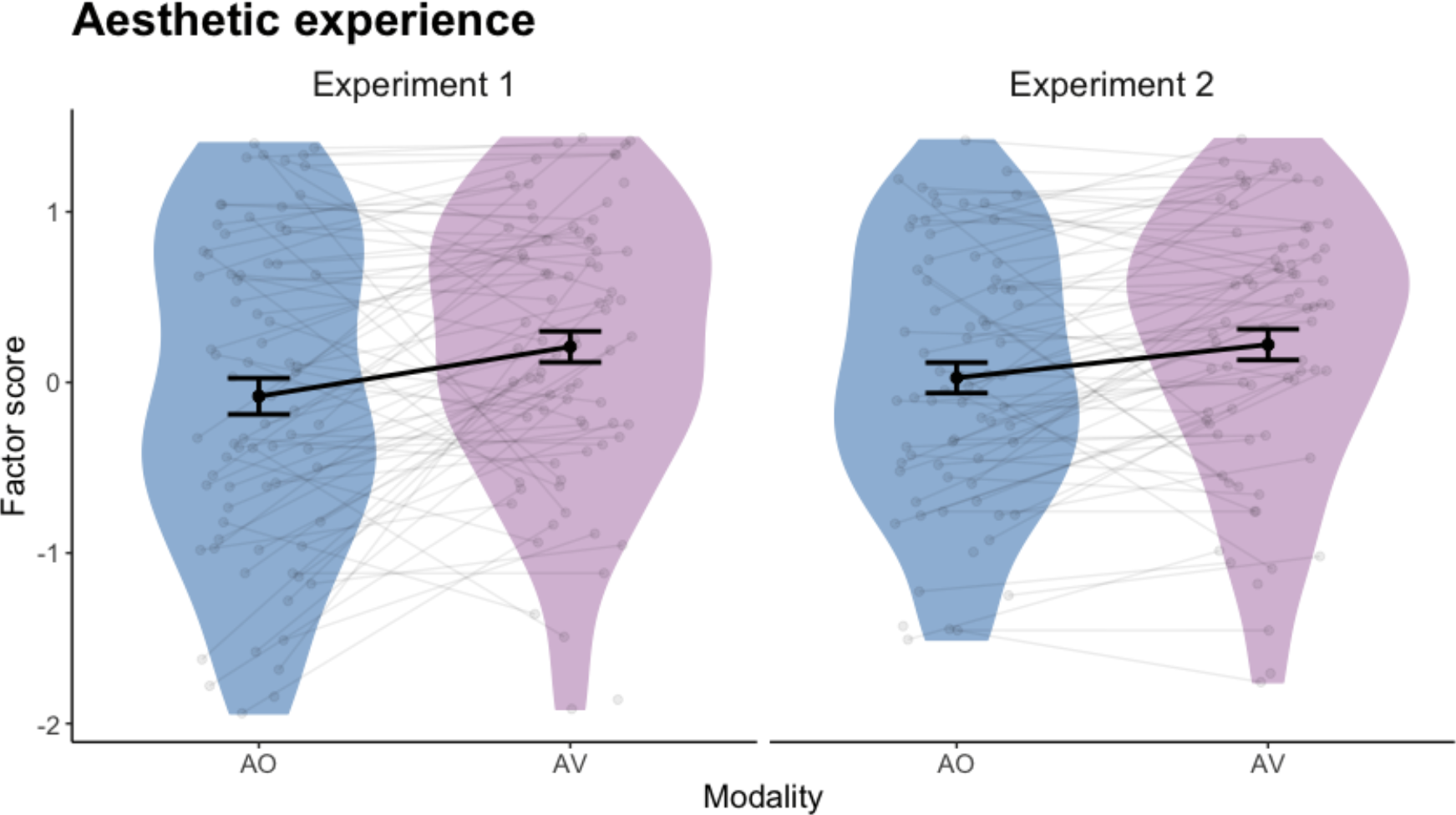
Aesthetic experience factor scores (which had high item loadings of liking, liking interpretation and absorption, see Table 1) as a function of modality (Audio Only (AO) is blue and Audiovisual (AV) is purple). The left panel shows results for Experiment 1, while the right panel shows results for Experiment 2. Each point represents factor scores for each participant and each piece.

### 3.3 Discussion

Experiment 1 tested whether participants had higher AE in the audio-only (AO) or audiovisual (AV) piano performances in a naturalistic concert setting. We confirmed that the measurements did not disturb participants and the findings show that AE increased in the AV compared to AO condition. These results support prior experimental laboratory results that showed liking and appreciation of expressivity are increased in AV conditions (Platz & Kopiez, 2012). We confirm that these results can be extended in a more naturalistic setting. One study that compared emotional differences between modalities in a naturalistic context, found higher wonder ratings but lower boredom ratings in live AV performances of music (Coutinho & Scherer, 2017). Our results likewise fit and extend this work, showing that the preference (liking) and absorption of the AE is also higher in AV modality. As naturalistic environments allow less control, it is important that these findings are replicated.

## 4 Experiment 2

Previous studies aimed at gaining further insight into potential emotional differences between uni- and bimodal music performances by measuring physiological responses (Chapados & Levitin, 2008; Vuoskoski et al., 2016). However, so far results are inconsistent. In Experiment 2, we explored whether different modalities would affect peripheral physiological responses similarly to the behavioural responses of AE (Exp. 1), and whether peripheral signals might serve as an index of AE.

### 4.1 Method

#### 4.1.1 Participants

The study was approved by the Ethics Council of the Max Planck Society and in accordance with the declarations of Helsinki. Participants gave their written informed consent. Twenty-six participants in total attended either Concert 3 (N=14) or Concert 4 (N = 12). Experiment 2 in total included nine females (17 males), with a mean age of 51.64 years (SD = 15.41), who on average had 5.94 (SD = 8.13) years of music lessons and attended an average of 14 concerts per year (M = 13.62, SD = 19.70). Participant provided ratings on their perception being a musician (from 1 = does not apply, to 7 completely applies), and most participants selected 1 (N = 15) or 2 (N = 3), while less selected 3 (N = 0), 4 (N= 1), 5 (N = 4), 6 (N = 2), or 7 (N = 1). All had either vocational training (*N* = 7) or a college/university degree (*N* = 19). Wilcoxon tests showed no significant differences between participants in Concert 3 and Concert 4 in terms of age (*p* = .72), years of music lessons (*p* = .14), and number of concerts attended in the last 12 months (*p* = 1.00). There was a significance in musician level between concerts (*p* = .039).

In assessing differences between the participant samples of the two Experiments, Experiment 1 had a significantly older audience on average (mean age in Experiment 1 = 58, Experiment 2 = 52, *p* = .041), but no significant differences for number of music lessons (*p* = .334), concert attendance in the last 12 months (*p* = .755), and musician level (*p* = .575).

Self-report data from all 26 participants were used in the analysis, while one physiological dataset from Concert 3 was lost due to technical problems (physiology: N = 25).

#### 4.1.2 Procedure

Participants were invited to arrive an hour before the concert, during which they were fitted with physiological equipment. All signals were collected with a portable recording device, ‘plux’ (https://plux.info/12-biosignalsplux), that continuously measured physiology across the duration of the concert at a 1000 Hz sampling rate. Respiration was measured via two respiration belts: one respiration belt was placed around the upper chest of the participant, and one respiration belt was placed around the lower belly. ECG, EMG, and EEG were collected using gelled self-adhesive disposable Ag/AgCl electrodes. Locations for the EMG, EEG, and ECG were prepared with peeling gel (under the left eyebrow and on left cheek for EEG, on the chest for ECG, and on the forehead for EEG). Three ECG electrodes were placed on the chest in a triangular arrangement; two as channels and one as the ground. Two facial muscles were recorded on the left side of participants’ faces; two electrodes were placed at the zygomaticus major (‘smiling’) muscle, and two electrodes were placed on the corrugator supercilii (‘frowning’) muscle, with a ground placed behind the left ear. EDA was collected via two electrodes placed on the middle phalanges of the non-dominant hand of participants. EEG activity from the frontal region was collected from three electrodes placed on the upper forehead, with a reference electrode placed in the middle of the forehead (in a similar location to an Fpz location in a conventional EEG cap), with additional two electrodes placed above the left and right eyebrows (in a similar position to Fp1 and Fp2 in a conventional EEG cap, respectively). EEG data are not reported in this paper.

#### 4.1.3 Factor analysis

We used the same items as in Experiment 1. Again, these item ratings were highly correlated (see accompanying code) and we chose to reduce these variables with a factor analysis. A Kaiser-Meyer-Olkin measure verified sampling adequacy (KMO = .609). All but one item had KMO values > .5; this one item (‘connection with co-listeners’) had a value of close to .5 (0.416). Correlations between items were large enough for a FA (Bartlett’s test of sphericity, *X*2(28) = 264.725, *p* < .001. Kaiser’s criterion of eigenvalues > 1 and a scree plot indicated a solution with one factor. Thus, a maximum likelihood factor analysis was conducted with one factor, which explained 24% of the variance. We took the scores from this factor and created a new variable. As we had similar loadings to Experiment 1, we also refer to this factor as the overall aesthetic experience (AE). In this factor, eleven outlier values exceeding ±3 Median Absolute Deviations (MAD, Leys et al., 2013) were removed from further analyses. In total, we had a total of 145 observations [(26 participants x 3 pieces x 2 modality conditions) - 11].

#### 4.1.4 Physiological pre-processing

Pre-processing of physiological signals (Experiment 2) was conducted in MATLAB (2019b, The Mathworks Inc, USA). Any missing data (gaps ranging from 5 - 53 ms long) were first linearly interpolated at the original sampling rate. Continuous data were then cut per piece. Using Ledalab (www.ledalab.de), skin conductance data were manually screened for artefacts (8% of data were rejected), downsampled to 20 Hz and separated into phasic (SCR) and tonic (SCL) components using Continuous Decomposition Analysis (Benedek & Kaernbach, 2010). Following previous literature, data were detrended to remove remaining long-term drifts (Omigie et al., 2021; cf. Salimpoor et al., 2009). Respiration, ECG, and EMG data were pre-processed using the Fieldtrip (Oostenveld et al., 2011) and biosig toolboxes in MATLAB (http://biosig.sourceforge.net/help/index.html). Manual screening of respiration data showed that the respiration signals obtained from the lower belly were stronger than those obtained from the upper chest; only data from the respiration belt around the lower belly were therefore used for further analysis. Respiration data were low-pass filtered at 2 Hz, ECG data were band-pass filtered between 0.6 and 20 Hz (Butterworth, 4^th^ order), and both demeaned. QRS peaks in the ECG signal were extracted using *nqrsdetect* function from biosignal, and peaks were found in respiration using custom functions. Computationally identified peaks were manually screened to ensure correct identification; any missing QRS peaks were manually added, while falsely identified QRS peaks were removed. Any ECG/respiration data that were too noisy for extraction of clear QRS/respiration peaks were rejected from further analysis (ECG = 14%, respiration = 7%). Differential timing of signal peaks – i.e., interbeat intervals (IBI, also known as RR-intervals) for ECG, and inter-breath intervals (IBrI) for respiration – were converted to beats per minute and interpolated at the original sampling rate to obtain a continuous respiration and heart rate. Heart rate variability measures were extracted using the *heartratevariability* function in biosig (http://biosig.sourceforge.net/). Normalised units of high frequency (HF, 0.15 – 0.4 Hz) power as well as the LF/HF ratio were taken into further analysis to reflect SNS and PNS activity (frequencies that adhere to the European Task Force recommendations (Malik, 1996). Electromyography (EMG) data for zygomaticus major (EMGZM) and corrugator supercilii (EMGCS) were band-pass filtered between 90 and 130 Hz and demeaned. We proceeded with the smoothed absolute value of the Hilbert transformed EMG signals.

Although there are questions as to what the most appropriate (central tendency) representation of physiological data is, we relied most closely on the methodology applied by Vuoskoski et al. (2016) to compare results. Therefore, the average of each (pre-processed) physiological measure was the main metric. As physiological responses change over time (i.e., they are non-stationary), and to gain a better representation (signal-to-noise ratio) of the responses across the course of each long piece, data for each piece were divided into piece sections that were driven by the musical structure (which were confirmed by a music theorist). Responses were averaged across these sections. Beethoven was split into nine, Messiaen into nine, and Bach into seven sections (see Supplementary Materials for more information). Overall, we were interested in eight physiological measures: averages of SCL, SCR, HR, HF power and LF/HF ratio, RR, as well as zygomaticus and corrugator activity, which we averaged per participant, modality, piece, and section. As with behavioural data, we removed outliers exceeding ±3 MAD. Total observations for each physiological measure after exclusion of noisy data and outliers were as follows: EMGCS = 1037, EMGZM = 1082, HR = 1073, HF = 1050, LF/HF ratio = 1041, RR = 1152, SCL = 1066, SCR = 910.

#### 4.1.5 Analysis

Statistical analysis for the AE scores obtained in Experiment 2 were conducted as described in Experiment 1. We also compared physiology between AO and AV modalities using LMMs (see General Methods, accompanying code). To determine if behavioural results were related to peripheral responses, we ran a LMM with aesthetic experience as the dependent variable and the eight peripheral measures (all of which were averaged across piece sections to represent rating per piece and scaled to be included in the same model) and condition as predictors. Random effect represented design-driven maximal were included: random intercepts were added for concert, piece, modality condition, and participant. Participants were nested within concerts, while participant, condition, and piece were considered as crossed effects. Variance Inflation Factors (VIF) were checked using the *car* package (Fox & Weisberg, 2019), confirming that VIFs were below 3.

### 4.2 Results

#### 4.2.1 Assessing naturalistic situations

We first assessed the extent to which the behavioural/physiological measurements disturbed the overall experience during the concert (i.e., for all pieces/conditions). Ratings suggested that measurements did not disrupt the concert experience, with a mean rating of 2.019 (SD = 1.416) and with 75% of ratings at 1 or 2 on the scale. Results are shown in Table 3. These results provide an important validation that physiological measurements can be used in the concert hall settings without impacting ecological validity.

**Table 3.**
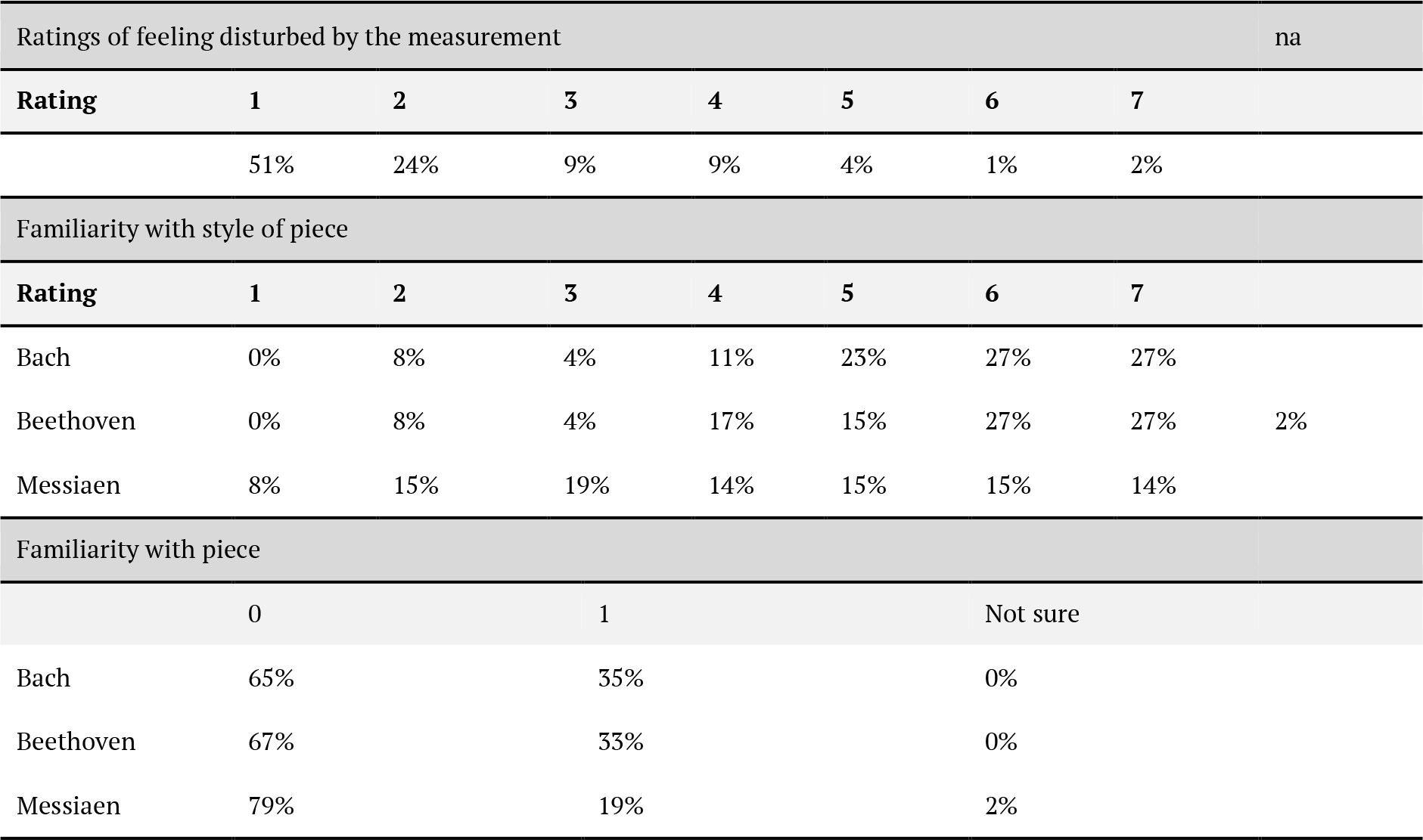
Ratings of feeling disturbed by the measurement and familiarity with style and specific piece in Experiment 2

**Table 4.** Linear mixed models for Aesthetic Experience factor scores between modality conditions.

**Aesthetic Experience (AE)**
| Predictors | Experiment 1 |  |  | Experiment 2 |  |  |
| --- | --- | --- | --- | --- | --- | --- |
|  | Estimates | CI | p | Estimates | CI | p |
| (Intercept) | -0.10 | -0.69 – 0.48 | 0.733 | 0.00 | -0.45 – 0.46 | 0.987 |
| cond [AV] | 0.29 | 0.05 – 0.52 | **0.018** | 0.22 | 0.01 – 0.43 | **0.040** |
| **Random Effects** |  |  |  |  |  |  |
| $\sigma^2$ | 0.55 | | | 0.40 | | |
| $\tau_{00}$ | 0.04 | id_n:concert | | 0.16 | id_n:concert | |
|  | 0.24 | piece |  | 0.08 | concert |  |
| ICC | 0.34 |  |  | 0.37 |  |  |
| N | 3 | piece |  | 2 | concert |  |
|  | 15 | id_n |  | 16 | id_n |  |
|  | 2 | concert |  |  |  |  |
| Observations | 153 |  |  | 145 |  |  |
| Marginal R<sup>2</sup> / Conditional R<sup>2</sup> | 0.024 / 0.355 |  |  | 0.019 / 0.383 |  |  |
| AIC | 371.967 |  |  | 324.565 |  |  |

#### 4.2.2 Piece familiarity

Similar to Experiment 1, ratings for familiarity of style were high for Bach (M = 5.385, SD = 1.484) and Beethoven (M = 5.333, SD = 1.532), but lower for Messiaen (M = 4.135, SD = 1.879). Approximately a third knew the Beethoven and Bach pieces, whereas only 19% knew the Messiaen piece.

#### 4.2.3 Aesthetic experience: Modality differences

For the behavioural AE results, LMMs showed modality was a significant predictor of AE (see Table 4) with AV scores significantly higher (M = .222, SE = 0.229, 95% CI [-2.07 2.52]) than AO scores (M = .003, SE = 0.229, 95% CI [-2.28 2.29], *t*(119) = -0.207, *p* = .041) (see Figure 2). Although the maximal model generated a singular fit error, it yielded the same estimate and significance, as well as a similar effect size and AIC to the simplified model that generated no error (see Supplementary Table 4). The modality effect was also confirmed by an ANOVA (*F*(1,25) = 6.832, *p* = .015). These results replicated the behavioural findings of Experiment 1.

#### 4.2.4 Physiological differences between modality

LMM results are presented in Table 5 (see also Figure 3). Modality condition was a significant predictor for LF/HF ratio, which represents SNS activation (higher arousal). Comparison of estimated marginal means indicated that this measure was higher in the AO than the AV condition (Table 6). This effect was consistent in the maximal models (see Supplementary Table 8) and confirmed by ANOVA (*F*(1,21) = 5.393, *p* = .030).

**Figure 3.**
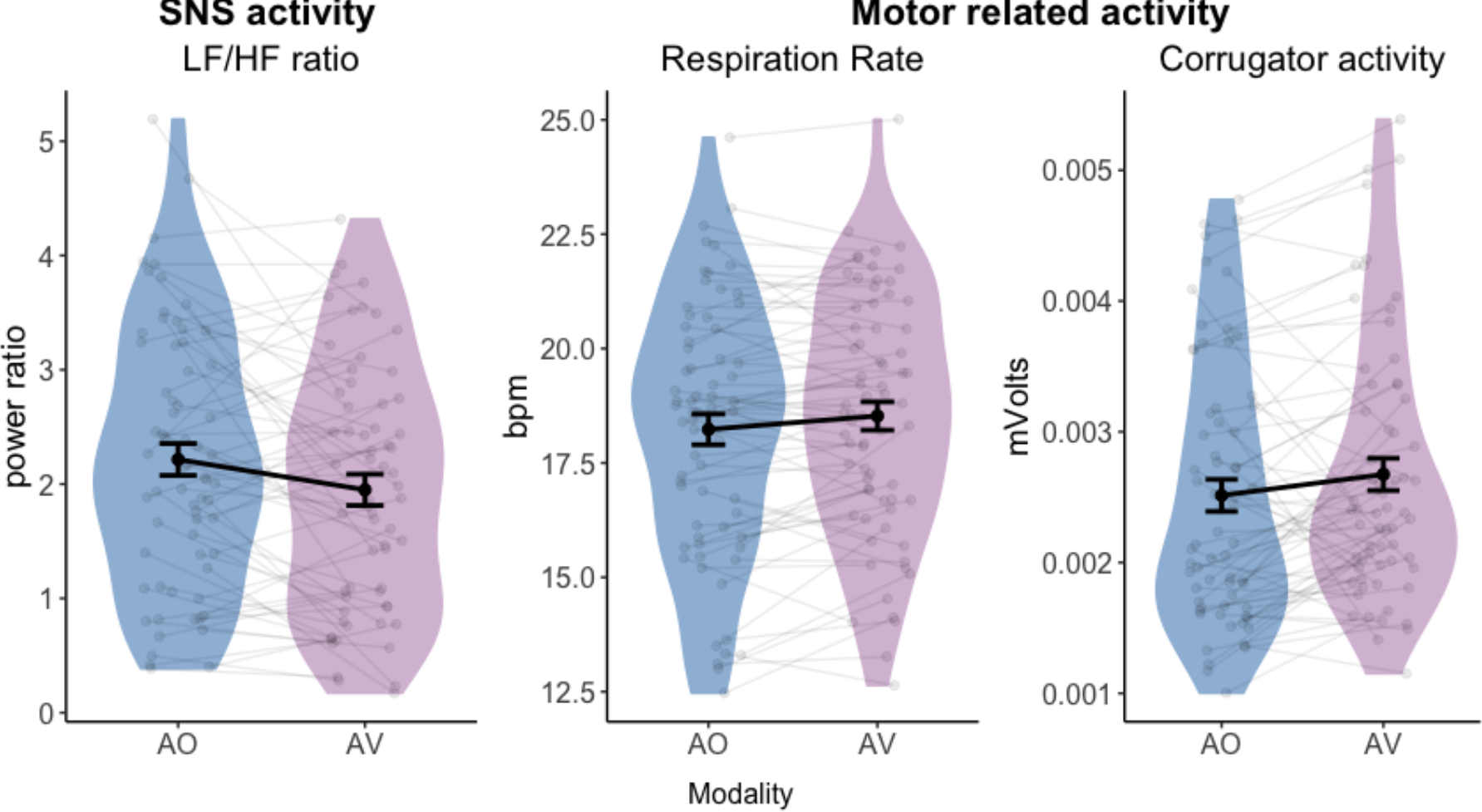
Physiological responses in each modality condition (AO: blue; AV: purple). Different panels represent different physiological measures; from left to right: LF/HF ratio, respiration rate (RR), and EMG activity of corrugator supercilii (frowning) muscle (Corrugator activity). Each point represents the physiological response value for each participant and each piece.

**Table 5.** Linear mixed models for physiological responses

| Physiological results |  |  |  |  |  |  |  |  |  |
| --- | --- | --- | --- | --- | --- | --- | --- | --- | --- |
| Predictors | EMGCS |  |  | EMGZM |  |  | HF |  |  |
|  | Estimates | CI | p | Estimates | CI | p | Estimates | CI | p |
| (Intercept) | 0.0025 | 0.0021 – 0.0029 | <0.001 | 0.0022 | 0.0019 – 0.0024 | <0.001 | 0.1384 | 0.1147 – 0.1621 | <0.001 |
| cond [AV] | 0.0002 | 0.0001 – 0.0003 | <0.001 | 0.0001 | -0.0000 – 0.0002 | 0.180 | 0.0044 | -0.0069 – 0.0157 | 0.447 |
| **Random Effects** |  |  |  |  |  |  |  |  |  |
| $\sigma^2$ | 0.00 | | | 0.00 | | | 0.00 | | |
| $\tau_{00}$ | 0.00 | section:piece | | 0.00 | section:piece | | 0.00 | section:piece | |
|  | 0.00 | id_n:concert |  | 0.00 | id_n:concert |  | 0.00 | id_n:concert |  |
| $\tau_{11}$ | | | | 0.00 | id_n:condAV | | 0.00 | id_n1:condAO | |
|  |  |  |  | 0.00 | id_n1:condAO |  | 0.00 | id_n2:condAV |  |
|  |  |  |  | 0.00 | id_n2:condAV |  |  |  |  |
| $\rho_{01}$ | | | | | | | | | |
| $\rho_{01}$ | | | | | | | | | |
| ICC | 0.65 |  |  | 0.50 |  |  | 0.47 |  |  |
| N | 9 | section |  | 9 | section |  | 9 | section |  |
|  | 3 | piece |  | 3 | piece |  | 3 | piece |  |
|  | 15 | id_n |  | 14 | id_n |  | 15 | id_n |  |
|  | 2 | concert |  | 2 | concert |  | 2 | concert |  |
| Observations | 1037 |  |  | 1082 |  |  | 1050 |  |  |
| Marginal R<sup>2</sup> / Conditional R<sup>2</sup> | 0.007 / 0.657 |  |  | 0.003 / 0.502 |  |  | 0.001 / 0.467 |  |  |

| Predictors | LF/HF ratio |  |  | HR |  |  | RR |  |  |
| --- | --- | --- | --- | --- | --- | --- | --- | --- | --- |
|  | Estimates | CI | p | Estimates | CI | p | Estimates | CI | p |
| (Intercept) | 2.19 | 1.78 – 2.61 | <0.001 | 62.00 | 54.96 – 69.04 | <0.001 | 18.31 | 17.17 – 19.45 | <0.001 |
| cond [AV] | -0.26 | -0.41 – -0.11 | 0.001 | -0.19 | -0.44 – 0.05 | 0.123 | 0.26 | 0.09 – 0.43 | 0.003 |
| **Random Effects** |  |  |  |  |  |  |  |  |  |
| $\sigma^2$ | 1.52 | | | 4.16 | | | 2.11 | | |
| $\tau_{00}$ | 0.00 | section:piece | | 0.11 | section:piece | | 0.30 | section:piece | |
|  | 0.95 | id_n:concert |  | 65.70 | id_n:concert |  | 6.71 | id_n:concert |  |
|  |  |  |  | 19.99 | concert |  | 0.08 | concert |  |
| ICC | 0.38 |  |  | 0.95 |  |  | 0.77 |  |  |
| N | 9 | section |  | 2 | concert |  | 2 | concert |  |
|  | 3 | piece |  | 9 | section |  | 9 | section |  |
|  | 15 | id_n |  | 3 | piece |  | 3 | piece |  |
|  | 2 | concert |  | 15 | id_n |  | 15 | id_n |  |
| Observations | 1041 |  |  | 1073 |  |  | 1152 |  |  |
| Marginal R<sup>2</sup> / Conditional R<sup>2</sup> | 0.007 / 0.389 |  |  | 0.000 / 0.954 |  |  | 0.002 / 0.771 |  |  |

| Predictors | SCR |  |  | SCL |  |  |
| --- | --- | --- | --- | --- | --- | --- |
|  | Estimates | CI | p | Estimates | CI | p |
| (Intercept) | -0.00 | -0.00 – 0.00 | 0.156 | 0.00 | -0.02 – 0.03 | 0.970 |
| cond [AV] | -0.00 | -0.00 – 0.00 | 0.754 | 0.00 | -0.01 – 0.02 | 0.764 |
| **Random Effects** |  |  |  |  |  |  |
| $\sigma^2$ | 0.00 | | | 0.01 | | |
| $\tau_{00}$ | 0.00 | section:piece | | 0.00 | section:piece | |
|  | 0.00 | id_n:concert |  |  |  |  |
| ICC | 0.25 |  |  | 0.21 |  |  |
| N | 9 | section |  | 9 | section |  |
|  | 3 | piece |  | 3 | piece |  |
|  | 14 | id_n |  |  |  |  |
|  | 2 | concert |  |  |  |  |
| Observations | 855 |  |  | 910 |  |  |
| Marginal R<sup>2</sup> / Conditional R<sup>2</sup> | 0.000 / 0.252 |  |  | 0.000 / 0.213 |  |  |

**Table 6.**
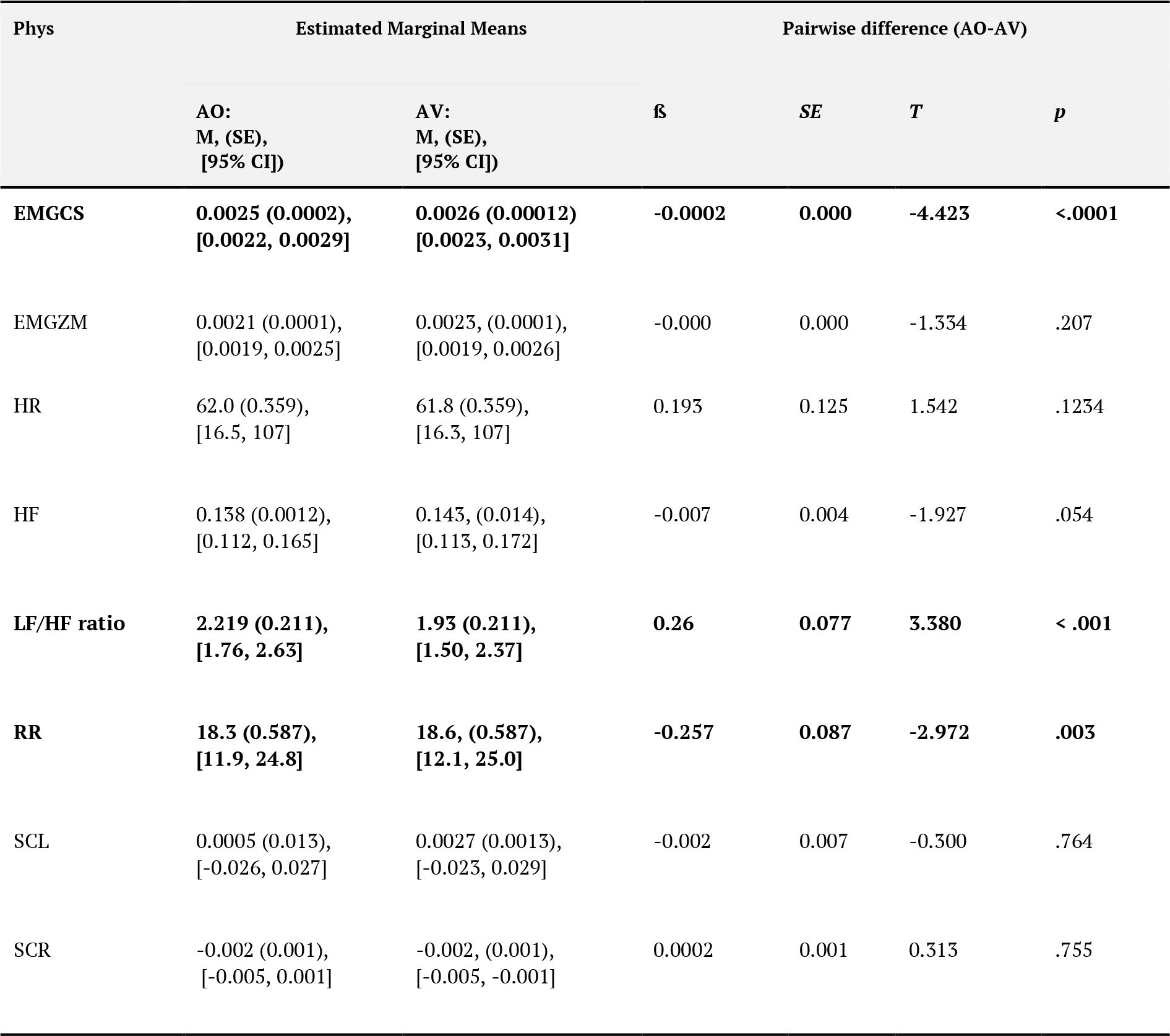
Results of Linear Mixed Models comparing Aesthetic Experience between AO and AV

Modality was a significant predictor for respiration rate (RR) and corrugator muscle activity (EMGCS), with a significant increase in the AV compared to AO condition (see Tables 5 and 6). However, in the maximal models that generated errors, the modality effect was not significant for EMGCS nor RR (see Supplementary Tables 5 and 10). Corresponding ANOVAs yielded insignificant results for RR (*F*(1,22) = 1.95, *p* = .177), though EMGCS was almost significant (*F*(1,21) = 3.679, *p* = .069). Due to the inconsistency of results between maximal models that generate errors and models with a simplified random structure that is free of errors, findings of EMGCS and RR are only cautiously interpreted.

#### 4.2.5 Peripheral measures that predict behaviour

In a model where AE was the dependent variable and all peripheral measures were predictors, zygomaticus activity (EMGZM) was a significant predictors of self-reported AE (see Table 7): increased smiling muscle activity was positively associated with AE.

**Table 7.** Model of physiology predicting AE self-reports

**LMM comparison physiology predicting Aesthetic Experience (AE)**
| Predictors | Maximal model |  |  |
| --- | --- | --- | --- |
|  | Estimates | CI | p |
| (Intercept) | 0.1940 | -0.1535 – 0.5415 | 0.270 |
| SCL | 0.0982 | -0.0737 – 0.2700 | 0.259 |
| SCR | -0.0108 | -0.1884 – 0.1668 | 0.904 |
| HR | -0.0500 | -0.2333 – 0.1333 | 0.589 |
| RR | 0.0628 | -0.0984 – 0.2241 | 0.440 |
| HFnu | -0.0007 | -0.2729 – 0.2715 | 0.996 |
| LFHFratio | -0.0351 | -0.3110 – 0.2409 | 0.801 |
| EMGCS | -0.0767 | -0.2508 – 0.0973 | 0.383 |
| EMGZM | 0.1828 | 0.0196 – 0.3460 | **0.029** |
| **Random Effects** |  |  |  |
| $\sigma^2$ | 0.49 | | |
| $\tau_{00}$ id_n:concert | 0.06 | | |
| $\tau_{00}$ piece | 0.00 | | |
| $\tau_{00}$ cond | 0.01 | | |
| $\tau_{00}$ concert | 0.03 | | |
| ICC | 0.18 |  |  |
| $N_{concert}$ | 2 | | |
| $N_{piece}$ | 3 | | |
| $N_{cond}$ | 2 | | |
| $N_{id\_n}$ | 13 | | |
| Observations | 94 |  |  |
| Marginal R<sup>2</sup> / Conditional R<sup>2</sup> | 0.065 / 0.234 |  |  |
| AIC | 256.674 |  |  |

### 4.3 Discussion

The main aims of Experiment 2 were to replicate the behavioural results of Experiment 1 and to gain further insight into peripheral physiological measures as a function of modality. Importantly, subjective ratings again showed that the measurement of physiological signals did not disturb participants.

As in Experiment 1, AE was significantly higher in the AV condition. We further tested whether peripheral responses between modality conditions. Compared to the AV condition, the AO condition evoked higher LF/HF ratio responses. These results support findings of (Vuoskoski et al., 2016), who reported higher physiological arousal in AO musical performances. On the other hand, respiration rate and corrugator muscle activity were higher in the AV condition. As both respiration and facial muscle activity are under voluntary muscle control, one interpretation is that viewing movements of the musician increased motor simulation. This is supported by research showing that viewing effortful movements increases respiration (Brown et al., 2013; Mulder et al., 2005; Paccalin & Jeannerod, 2000) and corrugator activity (de Morree & Marcora, 2010). However, inconsistencies occurred for RR and EMGCS in maximal LMMs compared to error-free LMMs. This model discrepancy suggests the modality effect in respiration and facial muscle activity needs to be complemented and confirmed by further studies with larger sample sizes.

When assessing if self-reported AE was predicted by physiological responses, AE was positively associated with zygomaticus activity. However, as increased zygomaticus activity has likewise been related to unpleasant experiences of (dissonant) music (Dellacherie et al., 2011; Merrill et al., 2021), we only cautiously attribute such facial muscle activity with positive AE.

## 5 General Discussion

The current experiments aimed to broaden our understanding of naturalistic concert experiences by testing whether (1) AV information enhances aesthetic experience (AE) in a more ecological setting and (2) peripheral physiological responses are higher in AO or AV modality. We also (3) assess the relationship between AE and peripheral physiological responses. We confirm that in both experiments, the measurement of self-report and physiology did not disturb the audiences, supporting the idea that a semi-experimental setting with naturalistic stimulus presentation can yield results of high ecological validity.

As there are several aspects that can make up an AE (Brattico & Pearce, 2013; Juslin, 2013; Schindler et al., 2017), questionnaire items related to certain aspects of an aesthetic experience were used. In both experiments, these items could be reduced to one factor in a factor analysis. Although the factor had slightly different loadings in the two experiments, three main items consistently loaded highly: absorption, liking, and liking of interpretation. Indeed, liking is a strong element of aesthetic experience both in philosophy (as the *evaluative dimension* of AE, Shusterman, 1997) and in empirical work (Brattico & Pearce, 2013). Preference of interpretation (e.g., how fast or expressive) has likewise been shown to play a strong role in AE. For example, observers prefer an expressive – compared to a non-expressive – interpretation of dance (Christensen et al., 2021). Similarly, dance choreography performed with more varied velocities was rated as more aesthetically pleasing compared to when it is performed with a more uniform velocity (Orlandi et al., 2020). Absorption has also shown to be an important factor in mediating aesthetic experience (Brattico & Pearce, 2013) and can even be indexed by peripheral measures, such as microsaccades (Lange et al., 2017). As these items have a strong connection to AE, it seemed appropriate to refer to this factor as such. Further, the fact that all of these items were correlated with each other and captured well by one factor, corroborates previous research that an aesthetic experience comprises many aspects (Brattico & Pearce, 2013; Merrill et al., 2021) and supports the use of dimensionality reduction techniques which trade specificity in favour of a more holistic AE measure.

Both Experiment 1 and 2 consistently showed that AE increases more in the AV than AO modality consistently across models and ANOVAs. Previous laboratory work has revealed that visual information carries several cues of musical expression (Davidson, 1993; Luck et al., 2010), quality (Tsay, 2013; Waddell & Williamon, 2017) and emotion (Dahl & Friberg, 2007; Van Zijl & Luck, 2013), which enhances aesthetic appreciation (Platz & Kopiez, 2012). Though these findings show that AE was significantly higher in AV than AO music performances, the effect size (just under 0.1) was relatively small (Cohen, 1988), likely due to the small sample size. Nonetheless, the overall model effect size (0.3 – 0.4) is considered medium (Cohen, 1988).

The current results extend the effect of modality influencing musical appreciation in a naturalistic performance setting. Similar work in a concert setting found that the live AV condition had increased wonder and decreased boredom (Coutinho & Scherer, 2017). However, their main focus was on emotion; we extend their findings to the preference (liking, liking of the interpretation) aspect of AE. We emphasise the importance of conducting AE research in a naturalistic performance setting, as it is more likely to elicit stronger and more realistic responses (Gabrielsson & Wik, 2003; Lamont, 2011). Of note is that results found in laboratory settings are not always replicated in more naturalistic settings. For example, previous laboratory studies have demonstrated that body movement (Platz & Kopiez, 2012) and familiarity (see North & Hargreaves, 2010) increase appreciation of music, even though the latter component has an inverted U-relationship. However, these findings were not replicated in a field study that was conducted in a more realistic situation (busking) and using a dependent variable of appreciation (i.e., money rather than ratings, Anglada-Tort et al., 2019), suggesting that components of music performance influence music appreciation differently depending on the context. Overall, despite the fact that a naturalistic setting might allow less control, together with results from previous work (Coutinho & Scherer, 2017), we provide consistent support that audiovisual information enhances AE; a finding that likely generalises to more naturalistic human behaviour.

We further elucidated peripheral responses of AE in multimodal contexts (Experiment 2), as research to date is inconsistent (Chapados & Levitin, 2008; Vuoskoski et al., 2016). Based on the framework of Brattico et al., (2013), we assume AE is made up of perceptual, cognitive, affective, and aesthetic responses (e.g., liking). These components can be relatively well distinguished by self-reports and – to an extent – by different brain regions and event-related brain potentials, depending on their latency and polarity (e.g., early components are related to early sensory processes). However, changes in physiology/facial muscle activity have been related to all of these cognitive, affective, and aesthetic responses (e.g., Roy et al., 2009; Steinbeis et al., 2006; Salimpoor et al., 2009), depending on the design and control condition of the study. Some show physiological changes related to sensory (orienting response, e.g., Barry & Sokolov, 1993) and acoustic changes (e.g., Chuen et al., 2016), while others show this activity is related to aesthetic preference (e.g., Grewe et al., 2009; Salimpoor et al., 2009). In further understanding physiological responses, we draw on neural and behavioural evidence that gives better insight into what kind of AE-related processing might take place. On the one hand, responses related to sensory processing should be greater in the AO condition, due to less predictable sound onsets (Jessen & Kotz, 2011), as also shown by Vuoskoski et al. (2016). On the other hand, AV information conveys more emotion (Dahl & Friberg, 2007; Van Zijl & Luck, 2013); therefore, responses could also be higher in the AV condition, as shown in Chapados and Levitin (2008). Thus, we tested again whether physiological responses are higher in AO or AV.

We consistently found that the LF/HF ratio increased in the AO condition. As this measure represents increased SNS activation, this suggests that AO conditions increase physiological arousal, likely reflecting an increase in uncertainty of sound onsets when visual information is absent (Jessen & Kotz, 2011; Klucharev et al., 2003; Stekelenburg & Vroomen, 2007; van Wassenhove et al., 2005). This is in line with results from Vuoskoski et al. (2016), who found that AO evoked more physiological arousal (as shown by skin conductance) compared to AV musical performances. We also support findings by Richardson et al. (2020) who likewise found higher physiological arousal in audio-only, compared to video versions of narratives (e.g., *Games of Thrones* and *Pride and Prejudice*).

We also found partial support for the hypothesis that AV music performances lead to higher peripheral physiological responses than in AO performances. We state partial evidence, as design-driven LMMs differed from error-free ones. Simplified, error-free models revealed a significant modality effect for RR and EMGCS. Maximal models, which generated errors, did not. These differences could be attributed to the fact that removing the slopes to avoid singularity fit errors could have increased degrees of freedom and the possibility of Type 1 errors (Arnqvist, 2019). However, a model with a complex random-effects structure can lead to increased Type II error and lack of power (Barr, 2021; Matuschek et al., 2017). Thus, future studies with larger sample sizes are required to confirm this modality effect. As there is general consensus that error-free models are preferable (Barr et al., 2013), these models are reported. Nonetheless, we aim to be transparent; the reader is pointed to not only the Supplementary Materials, but also the code showing the maximal models and how models are simplified step by step. While only cautiously interpreting the modality effects in RR and EMGCS, we believe it is worth briefly discussing the results from error-free models.

RR was faster in the AV condition. ‘Frowning’ muscle (EMGCS) activity, which typically reflects negative valence (Bradley & Lang, 2000), also increased in the AV condition. The discrepancy between the increase in both frowning muscle activity and (generally positive) AE in the AV condition could be explained by the fact that higher aesthetic pleasure can also derive from perceiving negatively valenced musical expression and/or affective states (Eerola et al., 2018), such as being moved (Eerola et al., 2016). However, some question whether facial expressions reflect valence (Wingenbach et al., 2020) or affective states at all (Lewis, 2011; Matsumo, 1987). Thus, another possible interpretation is that observing the musician increased mimicry in the observers. Indeed, participants mimic observed facial expressions (Dimberg, 1982; Magnee et al., 2007). Additionally, viewing effortful movements increases respiration (Brown et al., 2013; Mulder et al., 2005; Paccalin & Jeannerod, 2000) and corrugator activity (de Morree & Marcora, 2010). Such motor mimicry likely extends to music performance. Motor activity increases when listening to music (Bangert et al., 2006; Grahn & Brett, 2007; Janata et al., 2012), especially in audiovisual performances (Chan et al., 2013; Griffiths & Reay, 2018). Indeed, sensorimotor embodied mechanisms related to motor mimicry have been proposed and shown to enhance AE (Brattico & Pearce, 2013; Cross, 2011; Freedberg & Gallese, 2007; Gallese & Freedberg, 2007). Thus, faster breathing and increased facial muscle activity in AV conditions may be a reflection of motor mimicry that occurs when viewing musicians’ movements. In sum, we provide partial evidence of a modality effect in RR and EMGCS, potentially reflecting motor mimicry.

Facial muscle activity was significantly associated with AE. The zygomaticus (‘smiling’) muscle activity was a significant predictor for AE scores. Increased zygomaticus activity was positively related to AE, supporting previous work showing that zygomaticus activity was higher for pleasant music (Fuentes-Sánchez et al., 2022), liked positive music (Witvliet & Vrana, 2007), positively evaluated art (Gernot et al., 2018), and liked dance movements (Kirsch et al., 2016). This is further support for the embodied aesthetics theory, where sensorimotor embodied mechanisms might enhance AE (Brattico & Pearce, 2013; Cross, 2011; Freedberg & Gallese, 2007; Gallese & Freedberg, 2007). However, increased ‘smiling’ muscle activity has also been shown to increase in unpleasant (dissonant) music, suggesting that such activity might represent a grimace or ironic laughter (Dellacherie et al., 2011; Merrill et al., 2021). Therefore, it is vital to collect self-report data to support interpretations of physiological responses, rather than considering certain responses a direct index of a specific state, especially over a long period of time in such naturalistic settings.

LMMs show that LF/HF ratio were higher in AO, and tentative evidence suggests that respiration and muscle activity were higher in AV. These findings can be considered in conjunction with how much (in)voluntary control we have over them. As mentioned before, the heart is innervated by the ANS and made up of involuntary (cardiac) muscle. Voluntary skeletal muscles control EMG and (partly) respiration. On the one hand, due to the automatic nature of the heart, it seems plausible these might be more related to earlier (sensory) processes of an AE. On the other hand, the more voluntary peripheral measures seem to be related to the liking aspect of AE. Although we are cautious to attribute the increase of such measures as a direct index of aesthetic experience, the results point to the idea that the more voluntary the control of the peripheral measure, the more related it may be to later stages of the aesthetic processing, as outlined in Brattico and Pearce (2013).

One overall limitation of the current study is that although all versions were presented as part of a concert while participants were seated in the concert hall, AV was presented as a live version, while AO was presented as a playback. This was chosen to enhance ecological validity: people who listen to music in an AO version most likely listen to music as playback, while watching an AV version is more likely to be live (Sloboda et al., 2012). Indeed, this difference of visual information is also showed in Swarbrick et al. (2019), who similarly stated that AV performances are typically live. Although we do appreciate that tools and streaming platforms like YouTube, Digital Concert Hall of the Berliner Philharmoniker and MetOnDemand etc. have increased in popularity (especially with the COVID-19 pandemic) making audiovisual recording more popular, Belfi et al. (2021) found that felt pleasure did not differ between live and an audiovisual recording of that same performance. Therefore, it is likely that the live and playback differences do not play a strong role in influencing the current results. Future research might consider live audio-only playback of an offstage performer to fully mitigate this potential confound. Another limitation is that although the pieces were chosen to represent typical concert pieces (and a range of genres), they were not controlled for length. Nonetheless, length was a compromise when using naturalistic stimuli that heightened ecological validity. As we did not look at piece-specific differences, but rather average across sections of the pieces to examine the effect of condition, we did not consider this a confound in the current study. However, we note that effects driven by one piece may weigh our results more heavily than effects from the shorter pieces. Future research might consider choosing pieces of similar length, or at least similar lengths of sections. A further limitation is that we did not contrast visual only information with the other two conditions. This choice was a compromise to keep the within-in subject design time-manageable as well as to create a concert-like feel for the experiment.

## 6 Conclusion

Researchers are increasingly foregoing ultimate control for a more ecologically valid approach that enables participants to have more powerful aesthetic experiences. This study follows others that have moved more into the ‘wild’ to explore such naturalistic experiences (Chabin et al., 2022; Czepiel et al., 2021; Dotov & Trainor, 2021; Merrill et al., 2021; Swarbrick et al., 2019; Tervaniemi et al., 2021). The current findings show that a self-reported aesthetic experience significantly increases in audiovisual (compared to audio only) piano performances in the naturalistic setting of a concert hall.

Modality additionally influenced peripheral measures, revealing two main patterns. On the one hand, involuntary a physiological arousal response (heart rhythm reflecting SNS), was higher in the (less predictive) AO modality, likely reflecting more sensory processes. On the other hand, peripheral responses with more voluntary control (respiration, facial muscle activity) were higher in the AV modality, though due to inconsistencies in maximal/error-free models, these results should be interpreted with caution. The zygomaticus muscle was a significant predictor of self-reported AE. It could be that the involuntary-voluntary continuum of physiological responses is related to a sensory-affective continuum of AEs. We also suggest that visual information enhances motor mimicry (as shown by an increase in respiration and facial muscle activity), which is a mechanism that enhances AE (Cross et al., 2011; Freedberg & Gallese, 2007; Gallese & Freedberg, 2007; Kirsch et al., 2016). By exploring modality effects, we postulate that peripheral responses likely reflect sensory, sensorimotor, and affective responses that may culminate into an overall aesthetic experience (Brattico et al., 2013). However, we would like to emphasise that such peripheral responses alone cannot directly index AE; self-reports should support interpretations of peripheral physiological data. Nonetheless, the extent that physiological responses are simply sensory or reflect intertwined sensory and affective aspects of the aesthetic experience remains unclear. Further research, with larger sample sizes, should assess the robustness of the effects discussed here. To gain more insight, future research could bridge this gap by further exploring whether this involuntary-voluntary continuum reflects such sensory-aesthetic continuum and whether - and to what extent - there is an overlap of such systems.

## Acknowledgements

The authors thank Lea T. Fink for helping with music theoretical analysis, as well as the ArtLab team and assistants for help for the concerts. We thank Klaus Frieler for advice on statistical analysis and the Music Department of MPI EA in Frankfurt and Melanie Wald-Fuhrmann for discussions about the selection of musical pieces. We also thank the anonymous reviewers who provided insightful feedback for improving the manuscript.

## Author contribution

Using CRediT

Conceptualisation: CS

Methodology: CS

Investigation: CS, MS

Formal analysis: AC, SAK, MS, LKF

Writing - Original draft: AC

Visualisation: AC, LKF

Writing - review and editing: LKF, SAK, MS, CS

## Footnotes

1 The two main dimensions of emotion, according to the dimensional model of emotion(Russell, 1980). These terms reflect bipolar continuums: arousal ranging from calm to excitement, while valence varies from negative to positive emotional experience. Such peripheral responses have also been attributed to the discrete (basic) emotion theory, where SNS activation relates to happiness/fear, while PNS activation relates to calmness /sadness). For a more thorough discussion on emotion models, see for example (Barrett & Russell, 2015; Hamann, 2012).

