## Supplementary Materials for "Aesthetic and physiological effects of naturalistic multimodal music listening"

**Supplementary Table 1.** Correlations of musical features between concerts.

| Experiment 1 | | | | |
|---|---|---|---|---|
| Feature | Piece | R: C1-C2 (*p*) | R: C1-AO (*p*) | R: C2-AO (*p*) |
| RMS | Bach | 0.9185 ($p < .001$) | 0.8229 ($p < .001$) | 0.8539 ($p < .001$) |
| | Beet | 0.9777 ($p < .001$) | 0.9738 ($p < .001$) | 0.9537 ($p < .001$) |
| | Mess | 0.8813 ($p < .001$) | 0.8028 ($p < .001$) | 0.8196 ($p < .001$) |
| Spectral Centroid | Bach | 0.8160 ($p < .001$) | 0.7845 ($p < .001$) | 0.6093 ($p < .001$) |
| | Beet | 0.8568 ($p < .001$) | 0.7033 ($p < .001$) | 0.7400 ($p < .001$) |
| | Mess | 0.8869 ($p < .001$) | 0.8127 ($p < .001$) | 0.7751 ($p < .001$) |
| Tempo | Bach | 0.9195 ($p < .001$) | 0.8496 ($p < .001$) | 0.8316 ($p < .001$) |
| | Beet | 0.9328 ($p < .001$) | 0.9292 ($p < .001$) | 0.9136 ($p < .001$) |
| | Mess | 0.8861 ($p < .001$) | 0.8883 ($p < .001$) | 0.9283 ($p < .001$) |
| **Experiment 2** | | | | |
| Feature | Piece | R: C3-C4 (*p*) | R: C3-AO (*p*) | R: C4-AO (*p*) |
| RMS | Bach | 0.9498 ($p < .001$) | 0.8976 ($p < .001$) | 0.8969 ($p < .001$) |
| | Beet | 0.9898 ($p < .001$) | 0.9793($p < .001$) | 0.9844 ($p < .001$) |
| | Mess | 0.9574 ($p < .001$) | 0.9211 ($p < .001$) | 0.9194 ($p < .001$) |
| Spectral Centroid | Bach | 0.9539 ($p < .001$) | 0.6171 ($p < .001$) | 0.6415 ($p < .001$) |
| | Beet | 0.8931 ($p < .001$) | 0.8512 ($p < .001$) | 0.8200 ($p < .001$) |
| | Mess | 0.9222 ($p < .001$) | 0.9038 ($p < .001$) | 0.8805 ($p < .001$) |
| Tempo | Bach | 0.9337 ($p < .001$) | 0.9495 ($p < .001$) | 0.9583 ($p < .001$) |
| | Beet | 0.9713 ($p < .001$) | 0.9590 ($p < .001$) | 0.9706 ($p < .001$) |
| | Mess | 0.9815 ($p < .001$) | 0.9597 ($p < .001$) | 0.9706 ($p < .001$) |

**Supplementary Table 2.** Information and bar numbers of sections that pieces were divided into, driven by the musical structure

| Piece | Section | Bar corresponding to music | Bars corresponding to acoustic and physiological signal (i.e., considering repeats) | Approximate length in time (seconds) |
|---|---|---|---|---|
| Bach | 1 | Prelude: 1-16 | 1-16 | 44 |
| | 2 | Prelude: Repeat of bars 1-16 | 17-32 | 45 |
| | 3 | Prelude: 17-40 | 33-56 | 68 |
| | 4 | Prelude: 41 - 56 | 57-72 | 45 |
| | 5 | Prelude: Repeat of bars 17-40 | 73-96 | 68 |
| | 6 | Prelude: Repeat of bars 41-56 | 97-112 | 51 |
| | 7 | Fugue: 1-50 | 113-162 | 137 |
| Beethoven | 1 | 1-8 | 1-8 | 60 |
| | 2 | 9-16 | 9-16 | 46 |
| | 3 | 17-25 | 17-25 | 64 |
| | 4 | 26-29 | 26-29 | 27 |
| | 5 | 30-43 | 30-43 | 42 |
| | 6 | 44-55 | 44-55 | 42 |
| | 7 | 56-64 | 56-64 | 81 |
| | 8 | 65-75 | 65-63 | 64 |
| | 9 | 76-87 | 76-87 | 68 |
| Messiaen | 1 | 1-32 | 1-32 | 60 |
| | 2 | 33-40 | 33-40 | 41 |
| | 3 | 41-59 | 41-59 | 37 |
| | 4 | 60-131 | 60-131 | 142 |
| | 5 | 132-143 | 132-143 | 68 |
| | 6 | 144-174 | 144-174 | 43 |
| | 7 | 175-184 | 175-184 | 55 |
| | 8 | 185-216 | 185-216 | 64 |
| | 9 | 217-231 | 217-231 | 49 |

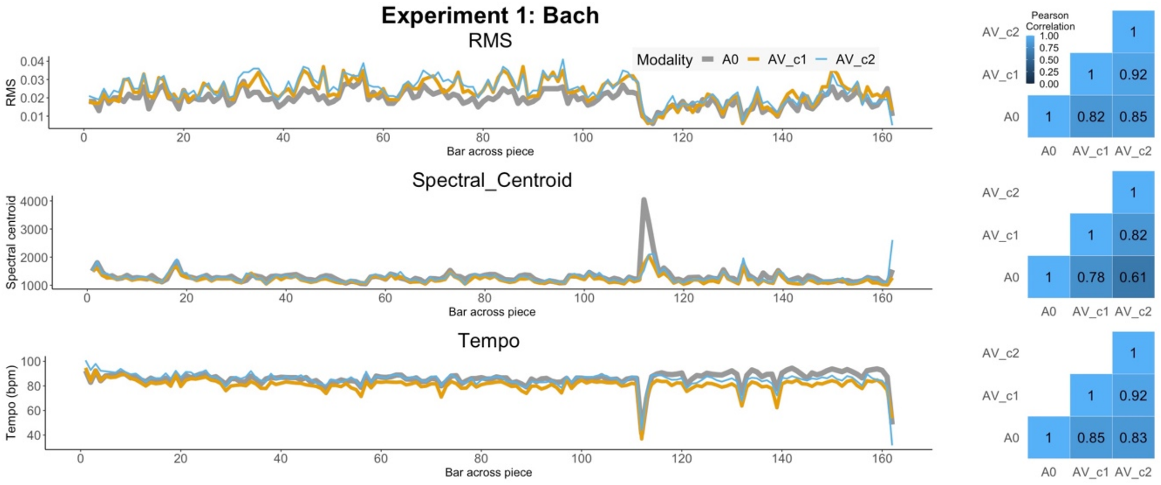


**Supplementary Figure 1.** Experiment 1, Bach piece: time series (left panels) and correlation matrices (right panels) of RMS, spectral centroid, and tempo (different rows) in AO and AV performances from concert 1 and 2.

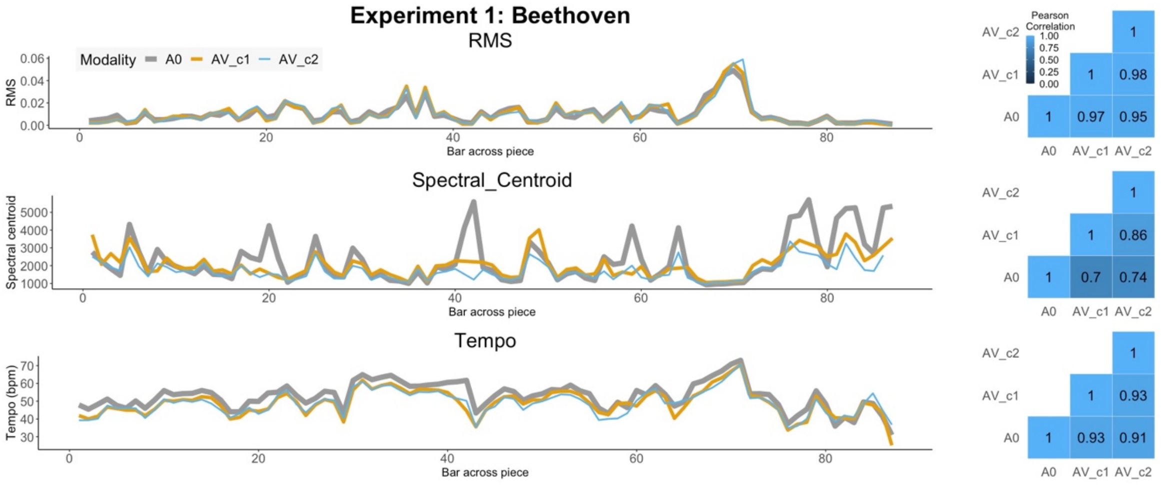


**Supplementary Figure 2.** Experiment 1, Beethoven piece: time series (left panels) and correlation matrices (right panels) of RMS, spectral centroid, and tempo (different rows) in AO and AV performances from concert 1 and 2.

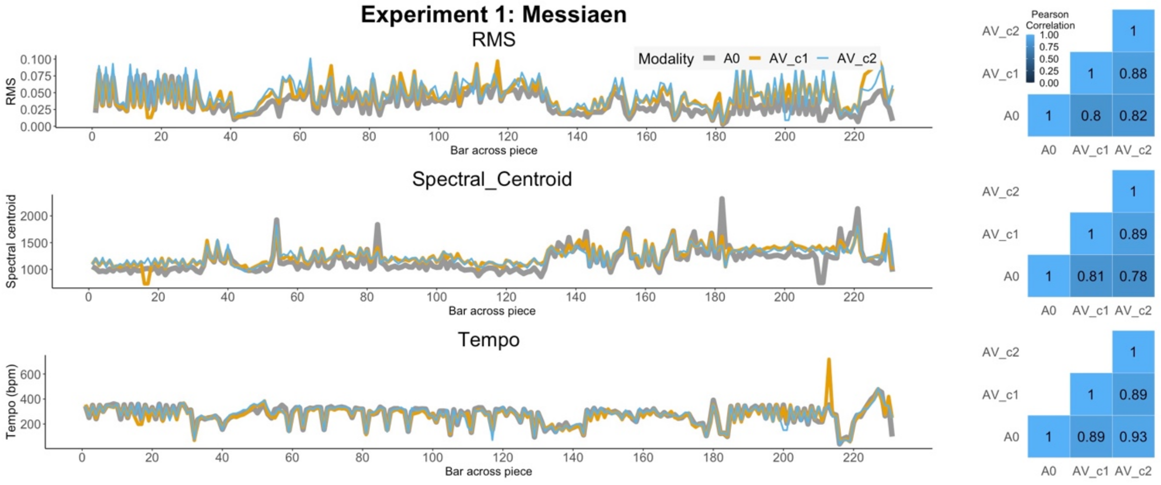


**Supplementary Figure 3.** Experiment 1, Messiaen piece: time series (left panels) and correlation matrices (right panels) of RMS, spectral centroid, and tempo (different rows) in AO and AV performances from concert 1 and 2.

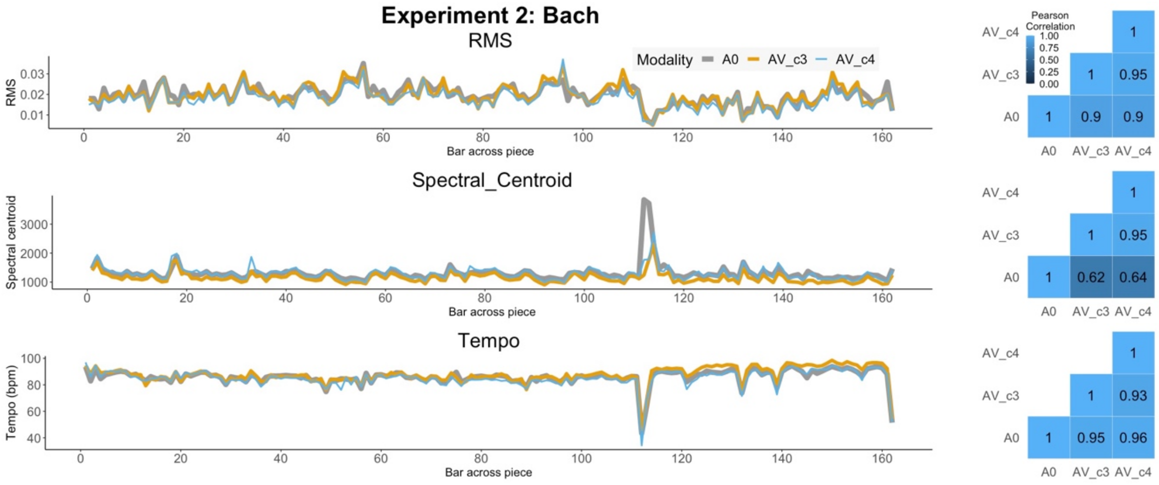


**Supplementary Figure 4.** Experiment 2, Bach piece: time series (left panels) and correlation matrices (right panels) of RMS, spectral centroid, and tempo (different rows) in AO and AV performances from concert 3 and 4.

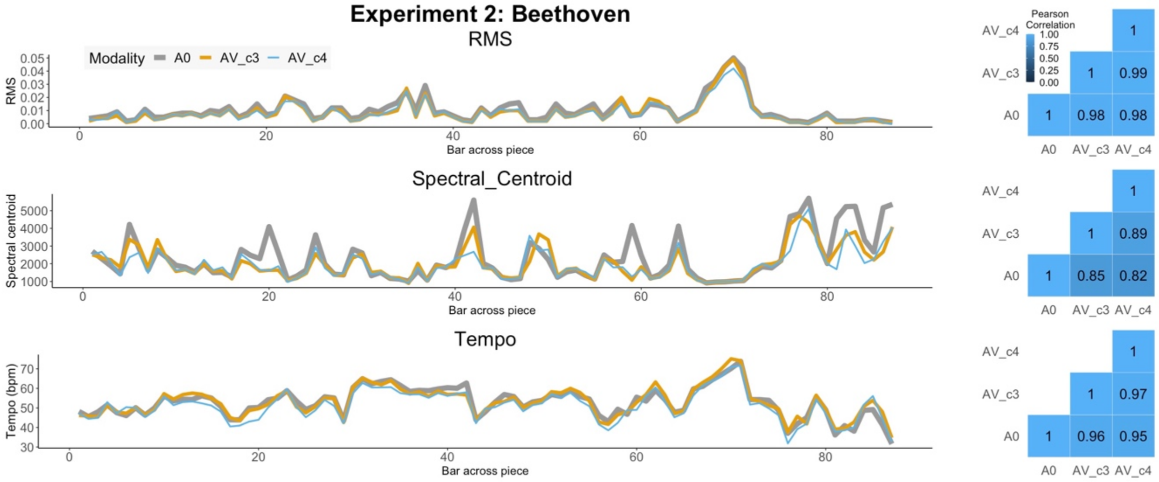


**Supplementary Figure 5.** Experiment 2, Beethoven piece: time series (left panels) and correlation matrices (right panels) of RMS, spectral centroid, and tempo (different rows) in AO and AV performances from concert 3 and 4.

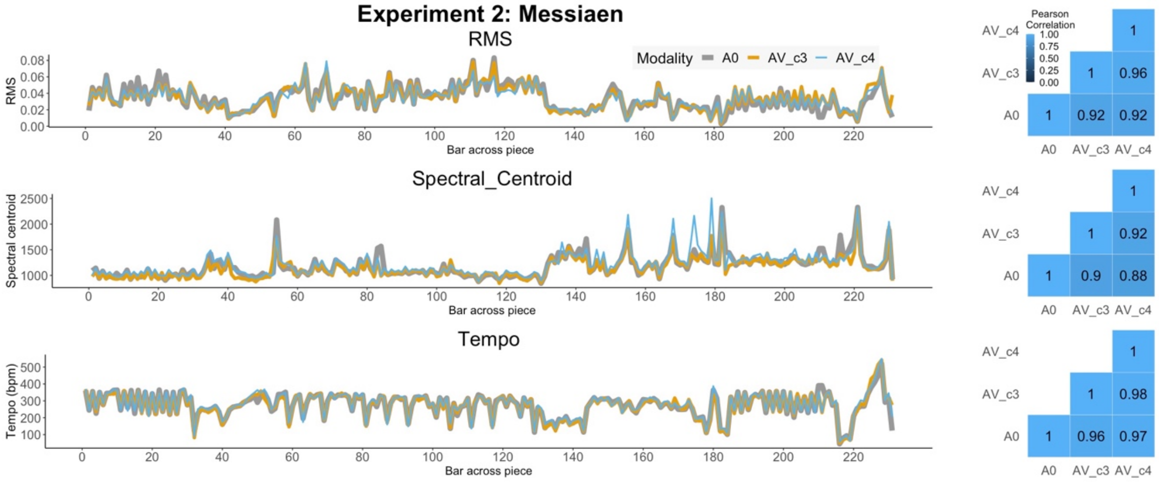


**Supplementary Figure 6.** Experiment 2, Messiaen piece: time series (left panels) and correlation matrices (right panels) of RMS, spectral centroid, and tempo (different rows) in AO and AV performances from concert 3 and 4.

**Supplementary Table 3.** Linear mixed models for aesthetic experience factor in Experiment 1: maximal and simplified random structure models, until no error is generated.

**LMM comparison for AE: Experiment 1**

| | **Maximal model (singularity error)** | | | **Constrained covariance parameters (singularity error)** | | | **Without slope (singularity error)** | | |
|---|---|---|---|---|---|---|---|---|---|
| *Predictors* | *Estimates* | *CI* | *p* | *Estimates* | *CI* | *p* | *Estimates* | *CI* | *p* |
| (Intercept) | -0.10 | -0.69 – 0.49 | 0.733 | -0.10 | -0.69 – 0.49 | 0.733 | -0.10 | -0.69 – 0.48 | 0.733 |
| cond [AV] | 0.29 | 0.05 – 0.52 | **0.018** | 0.29 | 0.05 – 0.52 | **0.018** | 0.29 | 0.05 – 0.52 | **0.018** |
| **Random Effects** | | | | | | | | | |
| $\sigma^2$ | 0.55 | | | 0.55 | | | 0.55 | | |
| $\tau_{00}$ | 0.04 $_{\text{id\_n:concert}}$ | | | 0.04 $_{\text{id\_n:concert}}$ | | | 0.04 $_{\text{id\_n:concert}}$ | | |
| | 0.24 $_{\text{piece}}$ | | | 0.24 $_{\text{piece}}$ | | | 0.24 $_{\text{piece}}$ | | |
| | 0.00 $_{\text{concert}}$ | | | 0.00 $_{\text{concert}}$ | | | 0.00 $_{\text{concert}}$ | | |
| $\tau_{11}$ | 0.00 $_{\text{id\_n.condAV}}$ | | | 0.00 $_{\text{id\_n.condAV}}$ | | | | | |
| | 0.00 $_{\text{id\_n1.condAO}}$ | | | 0.00 $_{\text{id\_n1.condAO}}$ | | | | | |
| | 0.00 $_{\text{id\_n2.condAV}}$ | | | 0.00 $_{\text{id\_n2.condAV}}$ | | | | | |
| $\rho_{01}$ | | | | | | | | | |
| $\rho_{01}$ | | | | | | | | | |
| N | 2 $_{\text{concert}}$ | | | 2 $_{\text{concert}}$ | | | 2 $_{\text{concert}}$ | | |
| | 3 $_{\text{piece}}$ | | | 3 $_{\text{piece}}$ | | | 3 $_{\text{piece}}$ | | |
| | 15 $_{\text{id\_n}}$ | | | 15 $_{\text{id\_n}}$ | | | 15 $_{\text{id\_n}}$ | | |
| Observations | 153 | | | 153 | | | 153 | | |
| Marginal $R^2$ / Conditional $R^2$ | 0.037 / NA | | | 0.037 / NA | | | 0.037 / NA | | |
| AIC | 379.967 | | | 379.967 | | | 373.967 | | |

**LMM comparison for AE: Experiment 1 (continued)**

| | **Without concert intercept (singularity error)** | | | **Without slope nor concert intercept (no error)** | | |
|---|---|---|---|---|---|---|
| *Predictors* | *Estimates* | *CI* | *p* | *Estimates* | *CI* | *p* |
| (Intercept) | -0.10 | -0.69 – 0.48 | 0.733 | -0.10 | -0.69 – 0.48 | 0.733 |
| cond [AV] | 0.29 | 0.05 – 0.52 | **0.018** | 0.29 | 0.05 – 0.52 | **0.018** |
| **Random Effects** | | | | | | |
| $\sigma^2$ | 0.55 | | | 0.55 | | |
| $\tau_{00}$ | 0.04 $_{\text{id\_n:concert}}$ | | | 0.04 $_{\text{id\_n:concert}}$ | | |
| | 0.24 $_{\text{piece}}$ | | | 0.24 $_{\text{piece}}$ | | |
| $\tau_{11}$ | 0.00 $_{\text{id\_n.condAV}}$ | | | | | |
| | 0.00 $_{\text{id\_n1.condAO}}$ | | | | | |
| | 0.00 $_{\text{id\_n2.condAV}}$ | | | | | |
| $\rho_{01}$ | | | | | | |
| $\rho_{01}$ | | | | | | |
| ICC | | | | 0.34 | | |
| N | 3 $_{\text{piece}}$ | | | 3 $_{\text{piece}}$ | | |
| | 15 $_{\text{id\_n}}$ | | | 15 $_{\text{id\_n}}$ | | |
| | 2 $_{\text{concert}}$ | | | 2 $_{\text{concert}}$ | | |
| Observations | 153 | | | 153 | | |
| Marginal $R^2$ / Conditional $R^2$ | 0.037 / NA | | | 0.024 / 0.355 | | |
| AIC | 377.967 | | | 371.967 | | |

**Supplementary Table 4.** Linear mixed models for aesthetic experience factor in Experiment 2: maximal and simplified random structure models, until no error is generated.

**LMM comparison for AE: Experiment 2**

| | Maximal model (singularity error) | | | Constrained covariance parameters (singularity error) | | | Without slope (singularity error) | | |
|---|---|---|---|---|---|---|---|---|---|
| *Predictors* | *Estimates* | *CI* | *p* | *Estimates* | *CI* | *p* | *Estimates* | *CI* | *p* |
| (Intercept) | 0.00 | -0.45 – 0.46 | 0.987 | 0.00 | -0.45 – 0.46 | 0.987 | 0.00 | -0.45 – 0.46 | 0.987 |
| cond [AV] | 0.22 | 0.01 – 0.43 | **0.040** | 0.22 | 0.01 – 0.43 | **0.040** | 0.22 | 0.01 – 0.43 | **0.040** |
| **Random Effects** | | | | | | | | | |
| $\sigma^2$ | 0.40 | | | 0.40 | | | 0.40 | | |
| $\tau_{00}$ | 0.15 $_{id\_n:concert}$ | | | 0.15 $_{id\_n:concert}$ | | | 0.16 $_{id\_n:concert}$ | | |
| | 0.00 $_{piece}$ | | | 0.00 $_{piece}$ | | | 0.00 $_{piece}$ | | |
| | 0.08 $_{concert}$ | | | 0.08 $_{concert}$ | | | 0.08 $_{concert}$ | | |
| $\tau_{11}$ | 0.00 $_{id\_n1.condAO}$ | | | 0.00 $_{id\_n1.condAO}$ | | | | | |
| | 0.00 $_{id\_n2.condAV}$ | | | 0.00 $_{id\_n2.condAV}$ | | | | | |
| $\rho_{01}$ | | | | | | | | | |
| $\rho_{01}$ | | | | | | | | | |
| N | 2 $_{concert}$ | | | 2 $_{concert}$ | | | 2 $_{concert}$ | | |
| | 3 $_{piece}$ | | | 3 $_{piece}$ | | | 3 $_{piece}$ | | |
| | 16 $_{id\_n}$ | | | 16 $_{id\_n}$ | | | 16 $_{id\_n}$ | | |
| Observations | 145 | | | 145 | | | 145 | | |
| Marginal $R^2$ / Conditional $R^2$ | 0.029 / NA | | | 0.029 / NA | | | 0.029 / NA | | |
| AIC | 332.563 | | | 332.563 | | | 326.565 | | |

**LMM comparison for AE: Experiment 2 (continued)**

| | Without piece intercept (singularity error) | | | Without slope nor piece intercept (no error) | | |
|---|---|---|---|---|---|---|
| *Predictors* | *Estimates* | *CI* | *p* | *Estimates* | *CI* | *p* |
| (Intercept) | 0.00 | -0.45 – 0.46 | 0.987 | 0.00 | -0.45 – 0.46 | 0.987 |
| cond [AV] | 0.22 | 0.01 – 0.43 | **0.040** | 0.22 | 0.01 – 0.43 | **0.040** |
| **Random Effects** | | | | | | |
| $\sigma^2$ | 0.40 | | | 0.40 | | |
| $\tau_{00}$ | 0.16 $_{id\_n:concert}$ | | | 0.16 $_{id\_n:concert}$ | | |
| | 0.08 $_{concert}$ | | | 0.08 $_{concert}$ | | |
| $\tau_{11}$ | 0.00 $_{id\_n.condAV}$ | | | | | |
| | 0.00 $_{id\_n1.condAO}$ | | | | | |
| | 0.00 $_{id\_n2.condAV}$ | | | | | |
| $\rho_{01}$ | | | | | | |
| $\rho_{01}$ | | | | | | |
| ICC | | | | 0.37 | | |
| N | 2 $_{concert}$ | | | 2 $_{concert}$ | | |
| | 16 $_{id\_n}$ | | | 16 $_{id\_n}$ | | |
| Observations | 145 | | | 145 | | |
| Marginal $R^2$ / Conditional $R^2$ | 0.029 / NA | | | 0.019 / 0.383 | | |
| AIC | 330.565 | | | 324.565 | | |

**Supplementary Table 5.** Linear mixed models for EMGCS (Corrugator facial muscle activity) in Experiment 2: maximal and simplified random structure models, until no error is generated.

**LMM comparison for EMGCS: Experiment 2**

| | Maximal model (singularity error) | | | Constrained covariance parameters (error) | | | Without slope (error) | | |
|---|---|---|---|---|---|---|---|---|---|
| *Predictors* | *Estimates* | *CI* | *p* | *Estimates* | *CI* | *p* | *Estimates* | *CI* | *p* |
| (Intercept) | 0.0025 | 0.0020 – 0.0030 | **<0.001** | 0.0025 | 0.0020 – 0.0030 | **<0.001** | 0.0025 | 0.0021 – 0.0029 | **<0.001** |
| cond [AV] | 0.0002 | -0.0000 – 0.0004 | 0.098 | 0.0002 | -0.0000 – 0.0004 | 0.098 | 0.0002 | 0.0001 – 0.0003 | **<0.001** |

**Random Effects**

| | Maximal model (singularity error) | Constrained covariance parameters (error) | Without slope (error) |
|---|---|---|---|
| $\sigma^2$ | 0.00 | 0.00 | 0.00 |
| $\tau_{00}$ | 0.00 section:piece | 0.00 section:piece | 0.00 section:piece |
| | 0.00 id_n:concert | 0.00 id_n:concert | 0.00 id_n:concert |
| | 0.00 concert | 0.00 concert | 0.00 concert |
| $\tau_{11}$ | 0.00 id_n.condAV | 0.00 id_n.condAV | |
| | 0.00 id_n1.condAO | 0.00 id_n1.condAO | |
| | 0.00 id_n2.condAV | 0.00 id_n2.condAV | |
| $\rho_{01}$ | | | |
| $\rho_{01}$ | | | |
| ICC | 0.61 | 0.61 | 0.65 |
| N | 2 concert | 2 concert | 2 concert |
| | 9 section | 9 section | 9 section |
| | 3 piece | 3 piece | 3 piece |
| | 15 id_n | 15 id_n | 15 id_n |
| Observations | 1037 | 1037 | 1037 |
| Marginal $R^2$ / Conditional $R^2$ | 0.010 / 0.617 | 0.010 / 0.617 | 0.007 / 0.657 |
| AIC | -12173.674 | -12173.674 | -12108.532 |

**LMM comparison for EMGCS: Experiment 2 (continued)**

| | Without concert intercept (error) | | | Without slope nor concert intercept (no error) | | |
|---|---|---|---|---|---|---|
| *Predictors* | *Estimates* | *CI* | *p* | *Estimates* | *CI* | *p* |
| (Intercept) | 0.0025 | 0.0020 – 0.0030 | **<0.001** | 0.0025 | 0.0021 – 0.0029 | **<0.001** |
| cond [AV] | 0.0002 | -0.0000 – 0.0004 | 0.098 | 0.0002 | 0.0001 – 0.0003 | **<0.001** |

**Random Effects**

| | Without concert intercept (error) | Without slope nor concert intercept (no error) |
|---|---|---|
| $\sigma^2$ | 0.00 | 0.00 |
| $\tau_{00}$ | 0.00 section:piece | 0.00 section:piece |
| | 0.00 id_n:concert | 0.00 id_n:concert |
| $\tau_{11}$ | 0.00 id_n.condAV | |
| | 0.00 id_n1.condAO | |
| | 0.00 id_n2.condAV | |
| $\rho_{01}$ | | |
| $\rho_{01}$ | | |
| ICC | 0.61 | 0.65 |
| N | 9 section | 9 section |
| | 3 piece | 3 piece |
| | 15 id_n | 15 id_n |
| | 2 concert | 2 concert |
| Observations | 1037 | 1037 |
| Marginal $R^2$ / Conditional $R^2$ | 0.010 / 0.618 | 0.007 / 0.657 |
| AIC | -12175.674 | -12110.532 |

**Supplementary Table 6.** Linear mixed models for EMGZM (Zygomaticus facial muscle activity) in Experiment 2: maximal and simplified random structure models, until no error is generated.

**LMM comparison for EMGZM: Experiment 2**

| | Maximal model (singularity error) | | | Constrained covariance parameters (error) | | |
|---|---|---|---|---|---|---|
| *Predictors* | *Estimates* | *CI* | *p* | *Estimates* | *CI* | *p* |
| (Intercept) | 0.0022 | 0.0019 – 0.0024 | **<0.001** | 0.0022 | 0.0019 – 0.0024 | **<0.001** |
| cond [AV] | 0.0001 | -0.0000 – 0.0002 | 0.180 | 0.0001 | -0.0000 – 0.0002 | 0.180 |
| **Random Effects** | | | | | | |
| $\sigma^2$ | 0.00 | | | 0.00 | | |
| $\tau_{00}$ | 0.00 section:piece | | | 0.00 section:piece | | |
| | 0.00 id_n:concert | | | 0.00 id_n:concert | | |
| | 0.00 concert | | | 0.00 concert | | |
| $\tau_{11}$ | 0.00 id_n.condAV | | | 0.00 id_n.condAV | | |
| | 0.00 id_n1.condAO | | | 0.00 id_n1.condAO | | |
| | 0.00 id_n2.condAV | | | 0.00 id_n2.condAV | | |
| $\rho_{01}$ | | | | | | |
| $\rho_{01}$ | | | | | | |
| N | 2 concert | | | 2 concert | | |
| | 9 section | | | 9 section | | |
| | 3 piece | | | 3 piece | | |
| | 14 id_n | | | 14 id_n | | |
| Observations | 1082 | | | 1082 | | |
| Marginal $R^2$ / Conditional $R^2$ | 0.006 / NA | | | 0.006 / NA | | |
| AIC | -12780.993 | | | -12780.993 | | |

**LMM comparison for EMGZM: Experiment 2 (continued)**

| | Without slope (error) | | | Without concert intercept (no error) | | |
|---|---|---|---|---|---|---|
| *Predictors* | *Estimates* | *CI* | *p* | *Estimates* | *CI* | *p* |
| (Intercept) | 0.0022 | 0.0019 – 0.0024 | **<0.001** | 0.0022 | 0.0019 – 0.0024 | **<0.001** |
| cond [AV] | 0.0001 | -0.0000 – 0.0001 | 0.052 | 0.0001 | -0.0000 – 0.0002 | 0.180 |
| **Random Effects** | | | | | | |
| $\sigma^2$ | 0.00 | | | 0.00 | | |
| $\tau_{00}$ | 0.00 section:piece | | | 0.00 section:piece | | |
| | 0.00 id_n:concert | | | 0.00 id_n:concert | | |
| | 0.00 concert | | | | | |
| $\tau_{11}$ | | | | 0.00 id_n.condAV | | |
| | | | | 0.00 id_n1.condAO | | |
| | | | | 0.00 id_n2.condAV | | |
| $\rho_{01}$ | | | | | | |
| $\rho_{01}$ | | | | | | |
| ICC | 0.50 | | | 0.50 | | |
| N | 2 concert | | | 9 section | | |
| | 9 section | | | 3 piece | | |
| | 3 piece | | | 14 id_n | | |
| | 14 id_n | | | 2 concert | | |
| Observations | 1082 | | | 1082 | | |
| Marginal $R^2$ / Conditional $R^2$ | 0.002 / 0.506 | | | 0.003 / 0.502 | | |
| AIC | -12771.767 | | | -12782.993 | | |

**Supplementary Table 7.** Linear mixed models for High Frequency power in heart rate variability (HF) in Experiment 2: maximal and simplified random structure models, until no error is generated.

**LMM comparison for HF: Experiment 2**

| | Maximal model (singularity error) | | | Constrained covariance parameters (error) | | |
|---|---|---|---|---|---|---|
| *Predictors* | *Estimates* | *CI* | *p* | *Estimates* | *CI* | *p* |
| (Intercept) | 0.1384 | 0.1147 – 0.1621 | **<0.001** | 0.1384 | 0.1147 – 0.1621 | **<0.001** |
| cond [AV] | 0.0044 | -0.0069 – 0.0157 | 0.447 | 0.0044 | -0.0069 – 0.0157 | 0.447 |

**Random Effects**

| | | |
|---|---|---|
| $\sigma^2$ | 0.00 | 0.00 |
| $\tau_{00}$ | 0.00 section:piece | 0.00 section:piece |
| | 0.00 id_n:concert | 0.00 id_n:concert |
| | 0.00 concert | 0.00 concert |
| $\tau_{11}$ | 0.00 id_n1.condAO | 0.00 id_n1.condAO |
| | 0.00 id_n2.condAV | 0.00 id_n2.condAV |
| $\rho_{01}$ | | |
| $\rho_{01}$ | | |
| ICC | 0.47 | 0.47 |
| N | 2 concert | 2 concert |
| | 9 section | 9 section |
| | 3 piece | 3 piece |
| | 15 id_n | 15 id_n |
| Observations | 1050 | 1050 |
| Marginal $R^2$ / Conditional $R^2$ | 0.001 / 0.467 | 0.001 / 0.467 |
| AIC | -2839.267 | -2839.267 |

**LMM comparison for HF: Experiment 2 (continued)**

| | Without slope (error) | | | Without concert intercept (no error) | | |
|---|---|---|---|---|---|---|
| *Predictors* | *Estimates* | *CI* | *p* | *Estimates* | *CI* | *p* |
| (Intercept) | 0.1394 | 0.1160 – 0.1628 | **<0.001** | 0.1384 | 0.1147 – 0.1621 | **<0.001** |
| cond [AV] | 0.0068 | -0.0004 – 0.0139 | 0.066 | 0.0044 | -0.0069 – 0.0157 | 0.447 |

**Random Effects**

| | | |
|---|---|---|
| $\sigma^2$ | 0.00 | 0.00 |
| $\tau_{00}$ | 0.00 section:piece | 0.00 section:piece |
| | 0.00 id_n:concert | 0.00 id_n:concert |
| | 0.00 concert | |
| $\tau_{11}$ | | 0.00 id_n1.condAO |
| | | 0.00 id_n2.condAV |
| $\rho_{01}$ | | |
| $\rho_{01}$ | | |
| ICC | | 0.47 |
| N | 2 concert | 9 section |
| | 9 section | 3 piece |
| | 3 piece | 15 id_n |
| | 15 id_n | 2 concert |
| Observations | 1050 | 1050 |
| Marginal $R^2$ / Conditional $R^2$ | 0.003 / NA | 0.001 / 0.467 |
| AIC | -2837.652 | -2841.267 |

**Supplementary Table 8.** Linear mixed models for Low Frequency / High Frequency ratio (LF/HF ratio) in heart rate variability in Experiment 2: maximal and simplified random structure models, until no error is generated.

**LMM comparison for LFHF ratio: Experiment 2**

| | Maximal model (singularity error) | | | Constrained covariance parameters (error) | | | Without slope (error) | | |
|---|---|---|---|---|---|---|---|---|---|
| *Predictors* | *Estimates* | *CI* | *p* | *Estimates* | *CI* | *p* | *Estimates* | *CI* | *p* |
| (Intercept) | 2.1904 | 1.7790 – 2.6017 | **<0.001** | 2.1904 | 1.7790 – 2.6017 | **<0.001** | 2.1926 | 1.7794 – 2.6059 | **<0.001** |
| cond [AV] | -0.2396 | -0.4304 – -0.0488 | **0.014** | -0.2396 | -0.4304 – -0.0488 | **0.014** | -0.2604 | -0.4116 – -0.1093 | **0.001** |
| **Random Effects** | | | | | | | | | |
| $\sigma^2$ | 1.51 | | | 1.51 | | | 1.52 | | |
| $\tau_{00}$ | 0.00 section:piece | | | 0.00 section:piece | | | 0.00 section:piece | | |
| | 0.94 id_n:concert | | | 0.94 id_n:concert | | | 0.95 id_n:concert | | |
| | 0.00 concert | | | 0.00 concert | | | 0.00 concert | | |
| $\tau_{11}$ | 0.00 id_n1.condAO | | | 0.00 id_n1.condAO | | | | | |
| | 0.04 id_n2.condAV | | | 0.04 id_n2.condAV | | | | | |
| $\rho_{01}$ | | | | | | | | | |
| $\rho_{01}$ | | | | | | | | | |
| ICC | 0.39 | | | 0.39 | | | 0.38 | | |
| N | 2 concert | | | 2 concert | | | 2 concert | | |
| | 9 section | | | 9 section | | | 9 section | | |
| | 3 piece | | | 3 piece | | | 3 piece | | |
| | 15 id_n | | | 15 id_n | | | 15 id_n | | |
| Observations | 1041 | | | 1041 | | | 1041 | | |
| Marginal $R^2$ / Conditional $R^2$ | 0.006 / 0.392 | | | 0.006 / 0.392 | | | 0.007 / 0.389 | | |
| AIC | 3489.670 | | | 3489.670 | | | 3485.161 | | |

**LMM comparison for LFHF ratio: Experiment 2 (continued)**

| | Without concert intercept (error) | | | Without slope nor concert intercept (no error) | | |
|---|---|---|---|---|---|---|
| *Predictors* | *Estimates* | *CI* | *p* | *Estimates* | *CI* | *p* |
| (Intercept) | 2.1903 | 1.7792 – 2.6015 | **<0.001** | 2.1926 | 1.7794 – 2.6059 | **<0.001** |
| cond [AV] | -0.2396 | -0.4304 – -0.0488 | **0.014** | -0.2604 | -0.4116 – -0.1093 | **0.001** |
| **Random Effects** | | | | | | |
| $\sigma^2$ | 1.51 | | | 1.52 | | |
| $\tau_{00}$ | 0.00 section:piece | | | 0.00 section:piece | | |
| | 0.94 id_n:concert | | | 0.95 id_n:concert | | |
| $\tau_{11}$ | 0.00 id_n1.condAO | | | | | |
| | 0.04 id_n2.condAV | | | | | |
| $\rho_{01}$ | | | | | | |
| $\rho_{01}$ | | | | | | |
| ICC | 0.39 | | | 0.38 | | |
| N | 9 section | | | 9 section | | |
| | 3 piece | | | 3 piece | | |
| | 15 id_n | | | 15 id_n | | |
| | 2 concert | | | 2 concert | | |
| Observations | 1041 | | | 1041 | | |
| Marginal $R^2$ / Conditional $R^2$ | 0.006 / 0.392 | | | 0.007 / 0.389 | | |
| AIC | 3487.670 | | | 3483.161 | | |

**Supplementary Table 9.** Linear mixed models for heart rate (HR) in Experiment 2: maximal and simplified random structure models, until no error is generated.

**LMM comparison for HR: Experiment 2**

| | Maximal model (error) | | | Constrained covariance parameters (error) | | | Without slope (no error) | | |
|---|---|---|---|---|---|---|---|---|---|
| *Predictors* | *Estimates* | *CI* | *p* | *Estimates* | *CI* | *p* | *Estimates* | *CI* | *p* |
| (Intercept) | 62.0143 | 54.3396 – 69.6890 | **<0.001** | 62.0143 | 54.3396 – 69.6890 | **<0.001** | 61.9969 | 54.9576 – 69.0361 | **<0.001** |
| cond [AV] | -0.2951 | -1.2997 – 0.7094 | 0.564 | -0.2951 | -1.2997 – 0.7094 | 0.564 | -0.1932 | -0.4390 – 0.0527 | 0.123 |

**Random Effects**

| | Maximal model (error) | Constrained covariance parameters (error) | Without slope (no error) |
|---|---|---|---|
| $\sigma^2$ | 3.46 | 3.46 | 4.16 |
| $\tau_{00}$ | $0.13_{\text{section:piece}}$ | $0.13_{\text{section:piece}}$ | $0.11_{\text{section:piece}}$ |
| | $65.05_{\text{id\_n:concert}}$ | $65.05_{\text{id\_n:concert}}$ | $65.70_{\text{id\_n:concert}}$ |
| | $24.91_{\text{concert}}$ | $24.91_{\text{concert}}$ | $19.99_{\text{concert}}$ |
| $\tau_{11}$ | $0.00_{\text{id\_n1.condAO}}$ | $0.00_{\text{id\_n1.condAO}}$ | |
| | $3.72_{\text{id\_n2.condAV}}$ | $3.72_{\text{id\_n2.condAV}}$ | |
| $\rho_{01}$ | | | |
| $\rho_{01}$ | | | |
| ICC | | | 0.95 |
| N | $2_{\text{concert}}$ | $2_{\text{concert}}$ | $2_{\text{concert}}$ |
| | $9_{\text{section}}$ | $9_{\text{section}}$ | $9_{\text{section}}$ |
| | $3_{\text{piece}}$ | $3_{\text{piece}}$ | $3_{\text{piece}}$ |
| | $15_{\text{id\_n}}$ | $15_{\text{id\_n}}$ | $15_{\text{id\_n}}$ |
| Observations | 1073 | 1073 | 1073 |
| Marginal $R^2$ / Conditional $R^2$ | 0.006 / NA | 0.006 / NA | 0.000 / 0.954 |
| AIC | 4613.406 | 4613.406 | 4754.982 |

**Supplementary Table 14.** Linear mixed models for respiration rate (RR) in Experiment 2: maximal and simplified random structure models, until no error is generated.

**LMM comparison for RR: Experiment 2**

| | Maximal model (singularity error) | | | Constrained covariance parameters (error) | | | Without slope (no error) | | |
|---|---|---|---|---|---|---|---|---|---|
| *Predictors* | *Estimates* | *CI* | *p* | *Estimates* | *CI* | *p* | *Estimates* | *CI* | *p* |
| (Intercept) | 18.2917 | 17.0021 – 19.5814 | **<0.001** | 18.2917 | 17.0021 – 19.5814 | **<0.001** | 18.3133 | 17.1729 – 19.4537 | **<0.001** |
| cond [AV] | 0.2274 | -0.1377 – 0.5925 | 0.222 | 0.2274 | -0.1377 – 0.5925 | 0.222 | 0.2570 | 0.0873 – 0.4266 | **0.003** |

**Random Effects**

| | Maximal model (singularity error) | Constrained covariance parameters (error) | Without slope (no error) |
|---|---|---|---|
| $\sigma^2$ | 2.03 | 2.03 | 2.11 |
| $\tau_{00}$ | $0.31_{\text{section:piece}}$ | $0.31_{\text{section:piece}}$ | $0.30_{\text{section:piece}}$ |
| | $4.71_{\text{id\_n:concert}}$ | $4.71_{\text{id\_n:concert}}$ | $6.71_{\text{id\_n:concert}}$ |
| | $0.00_{\text{concert}}$ | $0.00_{\text{concert}}$ | $0.08_{\text{concert}}$ |
| $\tau_{11}$ | $3.17_{\text{id\_n1.condAO}}$ | $3.17_{\text{id\_n1.condAO}}$ | |
| | $2.04_{\text{id\_n2.condAV}}$ | $2.04_{\text{id\_n2.condAV}}$ | |
| $\rho_{01}$ | | | |
| $\rho_{01}$ | | | |
| ICC | 0.75 | 0.75 | 0.77 |
| N | $2_{\text{concert}}$ | $2_{\text{concert}}$ | $2_{\text{concert}}$ |
| | $9_{\text{section}}$ | $9_{\text{section}}$ | $9_{\text{section}}$ |
| | $3_{\text{piece}}$ | $3_{\text{piece}}$ | $3_{\text{piece}}$ |
| | $15_{\text{id\_n}}$ | $15_{\text{id\_n}}$ | $15_{\text{id\_n}}$ |
| Observations | 1152 | 1152 | 1152 |
| Marginal $R^2$ / Conditional $R^2$ | 0.002 / 0.749 | 0.002 / 0.749 | 0.002 / 0.771 |
| AIC | 4294.608 | 4294.608 | 4312.895 |

**Supplementary Table 10.** Linear mixed models for skin conductance response (SCR) in Experiment 2: maximal and simplified random structure models, until no error is generated.

**LMM comparison for SCR: Experiment 2**

| | Maximal model (singularity error) | | | Constrained covariance parameters (error) | | | Without slope (error) | | |
|---|---|---|---|---|---|---|---|---|---|
| *Predictors* | *Estimates* | *CI* | *p* | *Estimates* | *CI* | *p* | *Estimates* | *CI* | *p* |
| (Intercept) | -0.0019 | -0.0045 – 0.0007 | 0.156 | -0.0019 | -0.0045 – 0.0007 | 0.156 | -0.0019 | -0.0045 – 0.0007 | 0.156 |
| cond [AV] | -0.0002 | -0.0016 – 0.0012 | 0.754 | -0.0002 | -0.0016 – 0.0012 | 0.754 | -0.0002 | -0.0016 – 0.0012 | 0.754 |
| **Random Effects** | | | | | | | | | |
| $\sigma^2$ | 0.00 | | | 0.00 | | | 0.00 | | |
| $\tau_{00}$ | 0.00 section:piece | | | 0.00 section:piece | | | 0.00 section:piece | | |
| | 0.00 id_n:concert | | | 0.00 id_n:concert | | | 0.00 id_n:concert | | |
| | 0.00 concert | | | 0.00 concert | | | 0.00 concert | | |
| $\tau_{11}$ | 0.00 id_n.condAV | | | 0.00 id_n.condAV | | | | | |
| | 0.00 id_n1.condAO | | | 0.00 id_n1.condAO | | | | | |
| | 0.00 id_n2.condAV | | | 0.00 id_n2.condAV | | | | | |
| $\rho_{01}$ | | | | | | | | | |
| $\rho_{01}$ | | | | | | | | | |
| N | 2 concert | | | 2 concert | | | 2 concert | | |
| | 9 section | | | 9 section | | | 9 section | | |
| | 3 piece | | | 3 piece | | | 3 piece | | |
| | 14 id_n | | | 14 id_n | | | 14 id_n | | |
| Observations | 855 | | | 855 | | | 855 | | |
| Marginal $R^2$ / Conditional $R^2$ | 0.000 / NA | | | 0.000 / NA | | | 0.000 / NA | | |
| AIC | -5275.751 | | | -5275.751 | | | -5281.751 | | |

**LMM comparison for SCR: Experiment 2 (continued)**

| | Without concert intercept (error) | | | Without slope nor concert intercept (no error) | | |
|---|---|---|---|---|---|---|
| *Predictors* | *Estimates* | *CI* | *p* | *Estimates* | *CI* | *p* |
| (Intercept) | -0.0019 | -0.0045 – 0.0007 | 0.156 | -0.0019 | -0.0045 – 0.0007 | 0.156 |
| cond [AV] | -0.0002 | -0.0016 – 0.0012 | 0.754 | -0.0002 | -0.0016 – 0.0012 | 0.754 |
| **Random Effects** | | | | | | |
| $\sigma^2$ | 0.00 | | | 0.00 | | |
| $\tau_{00}$ | 0.00 section:piece | | | 0.00 section:piece | | |
| | 0.00 id_n:concert | | | 0.00 id_n:concert | | |
| $\tau_{11}$ | 0.00 id_n.condAV | | | | | |
| | 0.00 id_n1.condAO | | | | | |
| | 0.00 id_n2.condAV | | | | | |
| $\rho_{01}$ | | | | | | |
| $\rho_{01}$ | | | | | | |
| ICC | | | | 0.25 | | |
| N | 9 section | | | 9 section | | |
| | 3 piece | | | 3 piece | | |
| | 14 id_n | | | 14 id_n | | |
| | 2 concert | | | 2 concert | | |
| Observations | 855 | | | 855 | | |
| Marginal $R^2$ / Conditional $R^2$ | 0.000 / NA | | | 0.000 / 0.252 | | |
| AIC | -5277.751 | | | -5283.751 | | |

**Supplementary Table 11.** Linear mixed models for skin conductance level (SCL) in Experiment 2: maximal and simplified random structure models, until no error is generated.

**LMM comparison for SCL: Experiment 2**

| | Maximal model (singularity error) | | | Constrained covariance parameters (error) | | | Without slope (error) | | |
|---|---|---|---|---|---|---|---|---|---|
| *Predictors* | *Estimates* | *CI* | *p* | *Estimates* | *CI* | *p* | *Estimates* | *CI* | *p* |
| (Intercept) | 0.0005 | -0.0247 – 0.0257 | 0.970 | 0.0005 | -0.0247 – 0.0257 | 0.970 | 0.0005 | -0.0247 – 0.0257 | 0.970 |
| cond [AV] | 0.0022 | -0.0124 – 0.0168 | 0.764 | 0.0022 | -0.0124 – 0.0168 | 0.764 | 0.0022 | -0.0124 – 0.0168 | 0.764 |

**Random Effects**

| | Maximal model (singularity error) | Constrained covariance parameters (error) | Without slope (error) |
|---|---|---|---|
| $\sigma^2$ | 0.01 | 0.01 | 0.01 |
| $\tau_{00}$ | 0.00 section:piece | 0.00 section:piece | 0.00 section:piece |
| | 0.00 id_n:concert | 0.00 id_n:concert | 0.00 id_n:concert |
| | 0.00 concert | 0.00 concert | 0.00 concert |
| $\tau_{11}$ | 0.00 id_n.condAV | 0.00 id_n.condAV | |
| | 0.00 id_n1.condAO | 0.00 id_n1.condAO | |
| | 0.00 id_n2.condAV | 0.00 id_n2.condAV | |
| $\rho_{01}$ | | | |
| $\rho_{01}$ | | | |
| N | 2 concert | 2 concert | 2 concert |
| | 9 section | 9 section | 9 section |
| | 3 piece | 3 piece | 3 piece |
| | 14 id_n | 14 id_n | 14 id_n |
| Observations | 910 | 910 | 910 |
| Marginal $R^2$ / Conditional $R^2$ | 0.000 / NA | 0.000 / NA | 0.000 / NA |
| AIC | -1312.587 | -1312.587 | -1318.587 |

**LMM comparison for SCL: Experiment 2 (continued)**

| | Without concert intercept (error) | | | Without slope nor concert intercept (error) | | | Without slope nor concert and id intercept (no error) | | |
|---|---|---|---|---|---|---|---|---|---|
| *Predictors* | *Estimates* | *CI* | *p* | *Estimates* | *CI* | *p* | *Estimates* | *CI* | *p* |
| (Intercept) | 0.0005 | -0.0247 – 0.0257 | 0.970 | 0.0005 | -0.0247 – 0.0257 | 0.970 | 0.0005 | -0.0247 – 0.0257 | 0.970 |
| cond [AV] | 0.0022 | -0.0124 – 0.0168 | 0.764 | 0.0022 | -0.0124 – 0.0168 | 0.764 | 0.0022 | -0.0124 – 0.0168 | 0.764 |

**Random Effects**

| | Without concert intercept (error) | Without slope nor concert intercept (error) | Without slope nor concert and id intercept (no error) |
|---|---|---|---|
| $\sigma^2$ | 0.01 | 0.01 | 0.01 |
| $\tau_{00}$ | 0.00 section:piece | 0.00 section:piece | 0.00 section:piece |
| | 0.00 id_n:concert | 0.00 id_n:concert | |
| $\tau_{11}$ | 0.00 id_n.condAV | | |
| | 0.00 id_n1.condAO | | |
| | 0.00 id_n2.condAV | | |
| $\rho_{01}$ | | | |
| $\rho_{01}$ | | | |
| ICC | | | 0.21 |
| N | 9 section | 9 section | 9 section |
| | 3 piece | 3 piece | 3 piece |
| | 14 id_n | 14 id_n | |
| | 2 concert | 2 concert | |
| Observations | 910 | 910 | 910 |
| Marginal $R^2$ / Conditional $R^2$ | 0.000 / NA | 0.000 / NA | 0.000 / 0.213 |
| AIC | -1314.587 | -1320.587 | -1322.587 |

**Questionnaire items (translated from German into English)**

0.1 Please state your age:

0.2 Please state your gender: 1. female; 2. male; 3. other

0.3 Please state your highest level of education:

1. Secondary school leaving certificate / Mittlere Reife
2. (Technical) Baccalaureate
3. Vocational training
4. College or university degree
5. not specified

0.4 How many years of instrumental lessons (including singing) have you had in your life? _____ years

0.5 Do you sing? (yes/no)

0.6 Do you play an instrument? (yes/no)

0.7 I would describe myself as a musician (1. do not agree to 7. Completely agree).

0.8 How many concerts/ live musical events have you attended within the last twelve months?

0.9 What kind of concerts/ live musical events do you attend most often? (rock/pop, classical, club/disco, jazz, contemporary, musical, opera, church, other).

1.1. Please answer the following questions

- 1.1.1 How much did you like the piece? (1-7)
- 1.1.2 How much did you like the interpretation? (1-7)
- 1.1.3 How familiar are you with this style of music that you have just heard? (1-7)
- 1.1.4 Do you know the piece? (yes/no)

1.2 To what extent do the following phrases apply to you?

- 1.2.1 I felt the need to move (1-7)
- 1.2.2 I tried to understand what was happening in the music (1-7)
- 1.2.3 I felt a connection to the musicians (1-7)
- 1.2.4 I was completely immersed in the music (1-7)
- 1.2.5 I felt connected to the other audience members (listeners) (1-7)
- 1.2.6 I was simply let the music affect me (passively receiving the music) (1-7)
- 1.2.7 I felt distracted by the measuring equipment (1-7)

1.3 Any other comments?
